# Role of desolvation on biomolecular liquid-liquid phase separation

**DOI:** 10.64898/2026.03.09.710469

**Authors:** Kai Zhang, Zhiyu Peng, Wenfei Li, Wei Wang

**Affiliations:** Department of Physics, National Laboratory of Solid State Microstructure, Nanjing University, Nanjing 210093, China; Wenzhou Key Laboratory of Biophysics, Wenzhou Institute, University of Chinese Academy of Sciences, Wenzhou, Zhejiang 325000, China; Jiangsu Key Laboratory for Cardiovascular Information and Health Engineering Medicine, Department of Cardiology, Nanjing Drum Tower Hospital, Medical School, Nanjing University, Nanjing 210093, P. R. China

**Author notes:** **For correspondence:** (WL); (WW).

## Abstract

Biomolecular condensates play essential roles in cellular organization and are implicated in diverse pathological processes. Their formation is driven by liquid-liquid phase separation (LLPS), a process that requires coordinated multistep desolvation of biomolecular chains and multivalent inter-chain interactions. Although coarse-grained (CG) models with implicit solvent are widely used to probe LLPS thermodynamics and kinetics, they typically neglect water-mediated desolvation effects, limiting their accuracy and mechanistic interpretability. Here, guided by all-atom simulations and experimental measurements, we develop a desolvation-aware implicit-solvent CG model by incorporating residue-level desolvation terms directly into the pairwise energy function and apply it to investigate LLPS of intrinsically disordered proteins. Incorporating these desolvation interactions reshapes the phase diagram, alleviating dense-phase overcompaction. Notably, we observe an approximately linear correlation between the temperature gap (simulation temperature relative to the critical point) and the extent of conformational expansion accompanying the dilute-to-dense phase transition, a result further supported by theoretical analysis. We also find that desolvation barriers slow early density-fluctuation growth and shorten transient kinetic arrest, whereas solvent-separated contact interactions exert the opposite effects. Both terms further modulate chain mobility within mature condensates through competing packing and energy-landscape effects. Together, this framework enables an efficient representation of desolvation in CG simulations and reveals how desolvation energetics shape both the thermodynamic landscape and kinetic properties of biomolecular LLPS.

## Introduction

Biomolecular condensates formed through liquid-liquid phase separation (LLPS) are increasingly recognized as fundamental regulators of diverse cellular processes, including stress responses ***Van Der Lee et al. (2014); Feric et al. (2016); Shin and Brangwynne (2017); Boeynaems et al. (2018); Burke et al. (2015); Biamonti and Vourc’h (2010); Riback et al. (2017***), cellular signaling ***Wippich et al. (2013); Su et al. (2016***), gene expression regulation ***Morimoto and Boerkoel (2013); Hnisz et al. (2017***), and chromatin organization ***Strom et al. (2017***). Dysregulation of condensate formation, in contrast, has been implicated in a wide range of human diseases, most notably cancer and numerous neurodegenerative disorders ***Molliex et al. (2015); Patel et al. (2015); Boeynaems et al. (2018); Conicella et al. (2016); Osterburg and Dötsch (2022***). Intrinsically disordered proteins (IDPs), which lack stable tertiary structures and often contain multivalent interaction motifs, are key molecular drivers of LLPS ***Molliex et al. (2015); Uversky et al. (2015); Zhou et al. (2018); Dignon et al. (2020); Elbaum-Garfinkle et al. (2015***). Understanding the molecular principles governing IDP-mediated LLPS is therefore essential for elucidating the mechanisms underlying both physiological condensate function and pathological phase transitions. A central yet incompletely understood aspect of LLPS is the multi-step desolvation of biomolecular chains as they transition from a homogeneously dispersed solution to a condensed phase. How desolvation energetics contribute to the thermodynamics and kinetics of LLPS remains a major open question.

Because experimentally characterizing the microscopic molecular events of LLPS at the sub-molecule level is still challenging for most experimental assays, a broad spectrum of computational approaches has been developed to characterize biomolecular phase separation. These range from mean-field ***Flory (1942); Huggins (1941***) and random-phase-approximation polymer theories ***Lin et al. (2017); Lin and Chan (2017); McCarty et al. (2019***) to diverse molecular simulation frameworks based on atomistic or coarse-grained (CG) force fields ***Dignon et al. (2018b***,a); ***Davtyan et al. (2012); De Jong et al. (2013); Baul et al. (2019***). Although all-atom molecular dynamics (MD) simulations can capture the microscopic processes of biomolecules with unprecedented temporal and spatial resolution, they become computationally prohibitive when applied to biomolecular phase separation involving a large number of protein chains. This challenge has motivated the development of CG models that reduce computational cost while retaining the essential physicochemical features governing phase behavior. These CG frameworks include early experiment-parameterized hydrophobicity-based models for IDPs ***Norgaard et al. (2008***), the sticker-and-spacer paradigm and its lattice Monte-Carlo implementation LASSI ***Harmon et al. (2017); Ruff et al. (2021); Choi et al. (2019***), the hydrophobicity-scale (HPS) family and its refinements ***Dignon et al. (2018b***,a); ***Regy et al. (2021); Dannenhoffer-Lafage and Best (2021***), and Mpipi, which explicitly incorporates cation-*π* interactions ***Joseph et al. (2021***). Advanced multiscale models have also been pioneered to predict the coupled phase behavior of IDPs and chromatin with remarkable accuracy ***Espinosa et al. (2020***). Higher-resolution or experimentally fine-tuned CG force fields (e.g., AWSEM-IDP ***Wu et al. (2018***), Martini3 ***Zhang et al. (2023); Souza et al. (2021***), COCOMO ***Valdes-Garcia et al. (2023); Jussupow et al. (2025***), AICG2+ ***Li et al. (2014); Bian et al. (2024***), and SIRAH ***Darré et al. (2015***)) further capture hydrogen bonding, secondary-structure propensity, and sequence-dependent packing. Notably, the maximum entropy optimized force field (MOFF) integrates maximum entropy biasing and energy landscape theory, successfully predicting the experimental dimensions and macroscopic phase separation of both ordered and disordered proteins ***Latham and Zhang (2019***, 2021). More recently, Tesei et al. introduced the CALVADOS model, which optimizes residue-specific interaction parameters within the HPS framework using SAXS and paramagnetic relaxation enhancement (PRE) data across a large IDP dataset ***Tesei et al. (2021); Tesei and Lindorff-Larsen (2022); Tesei et al. (2024***), thereby improving the accuracy and transferability of CG models for predicting IDP conformations and LLPS behavior.

Despite these advances, most residue-level CG models rely on implicit solvent representations, in which individual water molecules are not explicitly represented. This simplification can neglect the contribution of water molecules in mediating inter-residue interactions, which can be crucial for the thermodynamics and kinetics of biomolecular folding and phase transitions ***Cheung et al. (2002); Liu and Chan (2005); Karanicolas and Brooks III (2002); Tarus et al. (2006); Wu et al. (2011b***). In particular, the desolvation penalty associated with electrostatic and hydrophobic interactions exhibits marked differences between implicit and explicit solvent representations ***Salari and Chong (2012); Wu et al. (2011a***). More importantly, conventional implicit-solvent CG models used for LLPS usually do not directly account for the multi-step desolvation process that accompanies the transition from a dilute solution to a dense condensate. Recent theoretical and experimental studies highlight that water reorganization and solvent-mediated entropy changes are not merely passive consequences of LLPS but active driving forces that shape condensate stability, internal structure, and dynamics ***Mukherjee and Schäfer (2023); Mukherjee et al. (2024); Li and Hou (2023***). Related studies on hydrostatic pressure effects have further suggested that desolvation barriers and solvent-separated minima can help rationalize pressure-modulated LLPS behaviors, including the pressure-dependent reentrant phase separation of *α*-elastin ***Cinar et al. (2019***, 2018). Nevertheless, implicit-solvent CG models remain widely used because they strike an effective balance between computational efficiency and residue-level resolution. Incorporating desolvation effects into such implicit-solvent models thus represents a particularly promising strategy for enhancing their physical realism without sacrificing tractability in LLPS simulations. Notably, residue-level desolvation terms have already been successfully employed in protein folding studies ***Cheung et al. (2002); Liu and Chan (2005); Ferguson et al. (2009); Chen and Chan (2014); Zhang and Chan (2010); Karanicolas and Brooks III (2002); Gasic and Cheung (2020); Camilloni et al. (2016***). Within a structure-based modeling framework, Cheung et al. introduced a desolvation barrier separating the direct-contact minimum from the solvent-separated minimum in pairwise residue-residue potentials ***Cheung et al. (2002***). Simulations demonstrated that this minimalist desolvation model captures water-expulsion events during the final stages of folding. Chan and colleagues further showed that incorporating a desolvation barrier markedly enhances folding cooperativity ***Liu and Chan (2005); Ferguson et al. (2009); Chen and Chan (2014); Zhang and Chan (2010***). These previous successes provide a strong foundation for incorporating desolvation effects into coarse-grained models to more realistically investigate the critical contribution of desolvation to LLPS.

In this study, we advance the coarse-grained modeling of biomolecular LLPS by incorporating desolvation terms into the pairwise energy function of a CG model. By performing molecular simulations, we systematically investigate how desolvation influences phase behavior, thermodynamic properties, and the microscopic dynamics underlying condensate formation as well as the internal dynamics of the equilibrated dense phase. We further parameterize the desolvation terms using all-atom MD simulations and experimental *R*_*g*_ measurements, yielding a refined framework for investigating how generic desolvation features modulate the sequence-dependent phase behavior encoded by the underlying residue-level models.

## Results

### Effective representation of desolvation effects in coarse-grained models

In order to elucidate the role of water molecules in mediating inter-residue interactions, we first performed all-atom MD simulations with explicit solvent for a set of amino acid analogues and computed the potential of mean force (PMF) along inter-residue distances (*r*). Methane, methanol, acetate ion, ammonium ion, and methanamide were chosen to represent non-polar, polar, negatively charged, positively charged, and backbone-like residues, respectively. These amino acid analogues cover the key features of most amino acid types. The simulations were performed under ambient conditions (298 K and 1 atm), and the sampled snapshots were then analyzed to determine the inter-molecule distances and the corresponding PMF profiles. Additional simulation details are provided in the Methods section.

Figure 1A shows the PMF profile for the methane system. In addition to the deep potential well corresponding to the direct contact between two methane molecules (*r* ∼ 4 Å), there exists a shallow potential well at a larger distance (*r* ∼ 7 Å). This shallow minimum arises from transient, solvent-separated pseudo contacts between the solute residues, hereafter referred to as solvent-separated contact. To form a direct contact between the solutes, the water molecules mediating the solvent-separated contacts need to be excluded, which leads to a desolvation barrier between the two potential wells. Similar features have been reported in previous all-atom MD simulations ***Liu and Chan (2005***). The PMF profiles extracted from the simulations for other amino acid analogues exhibit the same overall shape (Figure 1B and ***Figure 1—figure Supplement 1***), but the positions of the direct-contact minimum (*r*_dc_), the solvent-separated minimum (*r*_ss_), and the desolvation barrier (*r*_b_) vary depending on the analogue molecules. The depths of the two potential wells (*ϵ* and *ϵ*_ss_) and the height of the desolvation barrier (*ϵ*_b_) likewise differ across analogue pairs.

**Figure 1.**
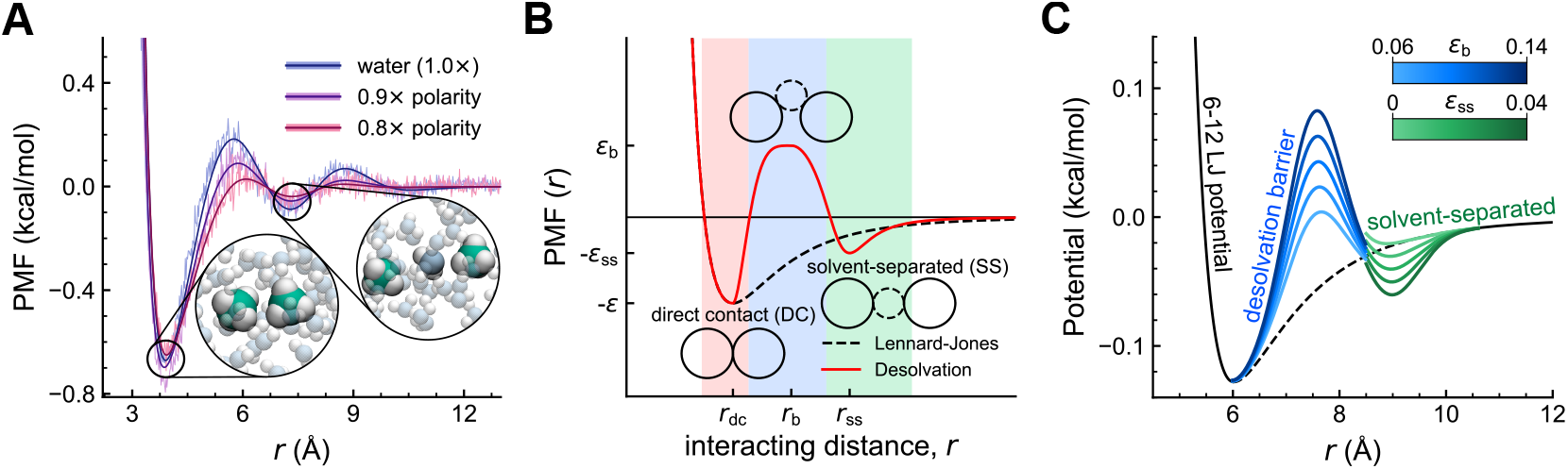
Desolvation-related PMF features of intermolecular interactions. (A) Potential of mean force along the intermolecular distances for the amino acid analogues from all-atom MD simulations with different solvent polarities. (B) Schematic diagram of desolvation effects. (C) Pairwise effective potential incorporating desolvation-inspired terms. Different curves correspond to different desolvation parameters.

The desolvation effect can also be modulated by solvent polarity and ionic conditions, which can be tuned experimentally by adding cosolvents and changing ionic strength, respectively. To examine the role of solvent polarity, we performed additional simulations of the methane system using water models with systematically varied polarity. For simplicity, the solvent polarity was tuned by uniformly scaling the partial charges of the hydrogen and oxygen atoms in the water molecules. This approach directly modulates the strength of solvent-solvent electrostatic interactions, which in turn dictates the stability of the hydration shell surrounding the solute. We compared the standard water model (1.0× charges) with reduced-polarity models (0.9× and 0.8× charges) (Figure 1A). The results reveal that a reduction in solvent polarity leads to a simultaneous decrease in the height of the desolvation barrier and the depth of the solvent-separated minimum. These observations underscore the importance of incorporating desolvation-related effective terms and exploring the effects of different desolvation strengths on the thermodynamics and kinetics of protein LLPS.

In this work, we introduce desolvation terms into the HPS model developed by Dignon et al. ***Dignon et al. (2018b***). In this framework, each residue is simplified to a single spherical bead. The HPS energy function includes a bonded term and a nonbonded term (see Materials and methods). The nonbonded term is described by the short-range pairwise interactions and long-range electrostatic interactions. To represent the desolvation features, we added two Gaussian terms centered at *r*_b_ and *r*_ss_, corresponding to the desolvation barrier and the solvent-separated minimum, respectively. The resulting nonbonded effective potential is given by

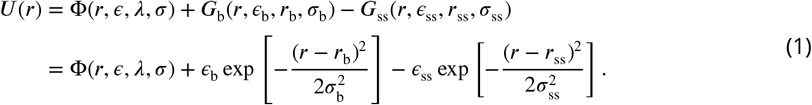

Here Φ(*r, ϵ, λ, σ*) denotes the nonbonded term in the original HPS model energy function ***Dignon et al. (2018b***). The parameters *λ* and *σ* correspond to the hydrophobicity scale values and effective bead sizes, respectively, for the 20 amino acids. The widths of the Gaussian terms were fixed at *σ*_b_ = *σ*_ss_ = 0.05 nm, while their centers were positioned at *r*_b_ = *r*_dc_+0.15 nm and *r*_ss_ = *r*_dc_+0.30 nm. These offsets approximately correspond to the radius and diameter of a water molecule, respectively.

We first employed a homopolymer protein model with a chain length of 50 residues to illustrate the desolvation effect. For each residue, we use the parameter set extracted by averaging over the IDPs in the DisProt dataset ***Aspromonte et al. (2024***), which gives an amino acid mass of *m*_residue_ = 110 amu (atomic mass unit), a hydrophobicity parameter *λ* = 0.640, and an effective bead size *σ* = 0.536 nm. The parameter *ϵ*_b_ controls the height of the desolvation barrier associated with the transition from solvent-separated to direct-contact configurations, whereas *ϵ*_ss_ determines the depth of the local minimum corresponding to a solvent-separated contact. Together, these parameters shape the desolvation-inspired effective potential and modulate the statistical balance between direct-contact and solvent-separated configurations. By tuning the parameter values of *ϵ*_ss_ and *ϵ*_b_, we generate energy functions with varying desolvation contributions (Figure 1C), enabling systematic investigation of how desolvation influences the phase behavior and thermodynamics of IDPs. Because the effective interaction parameters used in the model are temperature-independent, the absolute simulation temperature should not be directly interpreted as an experimental temperature. We therefore report all thermodynamic quantities, unless explicitly stated otherwise, in terms of a reduced temperature, defined as *T* ^∗^ = *k*_B_*T* /*ϵ*. Here, *ϵ* denotes the characteristic interaction energy scale of the system, which is set to *ϵ* = 0.2 kcal/mol following the convention of the HPS model ***Dignon et al. (2018b***). Accordingly, *T* ^∗^ should be interpreted primarily as a model temperature that controls the relative strength of thermal fluctuations, rather than as having a direct quantitative correspondence with experimental temperature.

### Thermodynamic impact of desolvation on the phase diagram

To capture the essential physics of IDP phase separation, we employed a coarse-grained model of 125 homopolymer chains using a slab simulation protocol (Methods). As a control, we first performed molecular dynamics simulations within the HPS model framework without considering the desolvation terms across a range of temperatures (Figure 2A–C). Representative snapshots clearly illustrate the transition from a homogeneous solution to a stable condensate as the temperature decreases (Figure 2B). We extracted time-averaged density profiles along the *z*-axis to identify coexisting dense and dilute phases and thereby constructed the binodal curve (Figure 2A, C), which reproduces phase behavior reported in previous molecular simulation studies ***Dignon et al. (2018b***). We then repeated the slab simulations using desolvation-inspired effective potentials with varying *ϵ*_b_ and *ϵ*_ss_ to investigate how desolvation reshapes the phase diagram and condensate structure.

**Figure 2.**
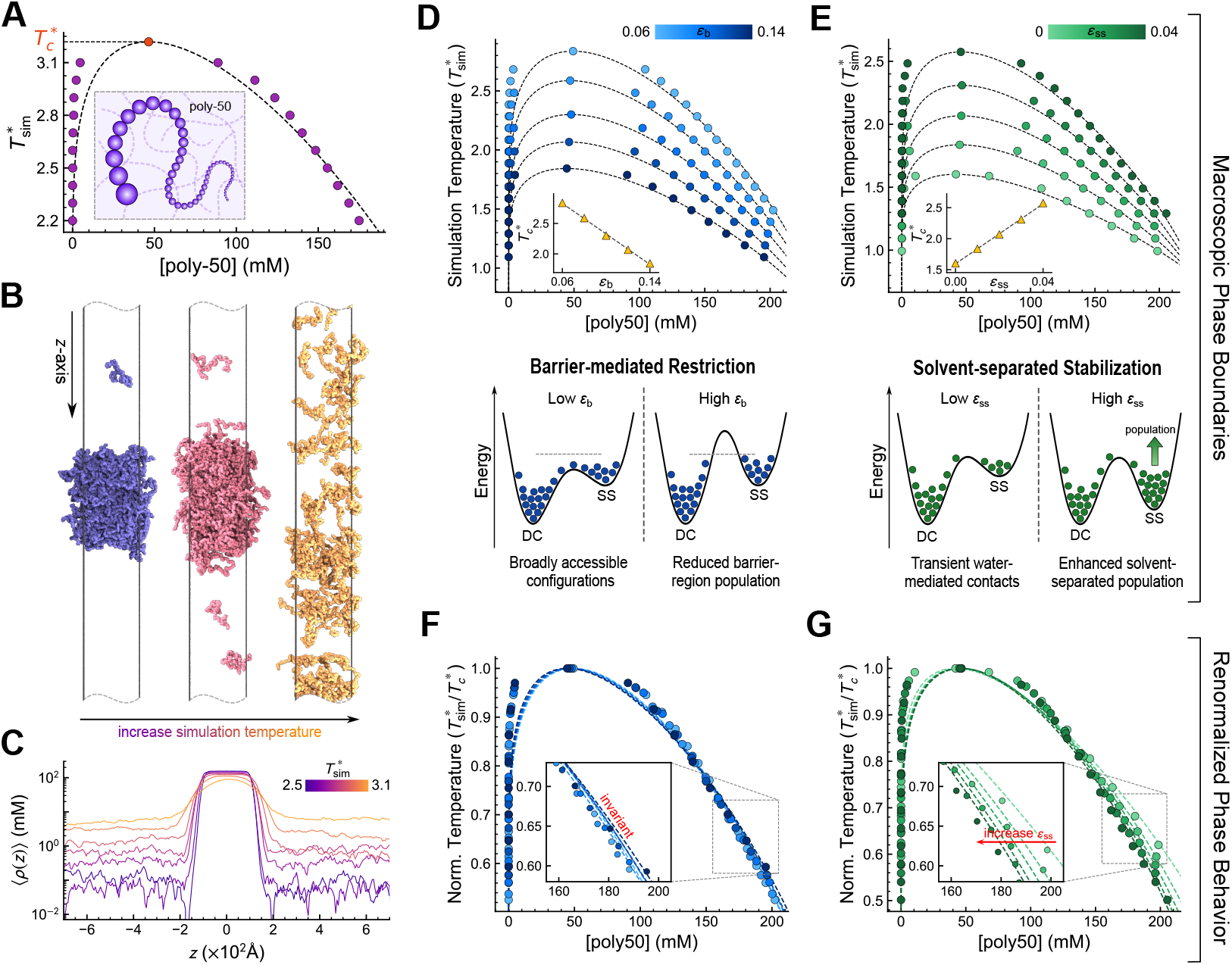
Thermodynamic regulation and microscopic mechanisms of desolvation-mediated phase separation. (A) Baseline phase diagram of the poly-50 system using the standard HPS model. (B) Representative simulation snapshots visualizing the transition from a stable condensate (*T* ^∗^ = 2.58) to a near-critical state (*T* ^∗^ = 2.98) and a homogeneous solution (*T* ^∗^ = 3.18). (C) Time-averaged density profiles along the *z*-axis identifying the coexisting dense and dilute phases. (D, E) Macroscopic phase boundaries under varying desolvation barrier heights *ϵ*_b_ (D) and solvent-separated potential depths *ϵ*_ss_ (E). Insets show the monotonic dependence of *T*_*c*_ on the respective parameters. The lower panels schematically illustrate how changes in *ϵ*_b_ and *ϵ*_ss_ alter the distribution of residue-pair configurations. The small circles indicate schematic populations of residue-pair configurations along the potential profile, with denser circles representing a higher population. (F, G) Renormalized phase behavior plotted against normalized temperature *T* ^∗^/*T*_*c*_ for varying *ϵ*_b_ (F) and varying *ϵ*_ss_ (G).

To elucidate how desolvation regulates the thermodynamics of LLPS, we systematically mapped phase diagrams under varying desolvation parameters (Figure 2D and E). Increasing the desolvation barrier *ϵ*_b_ monotonically lowers the critical temperature 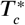 (Figure 2D), suggesting a reduced phase separation propensity. Analysis of residue-residue radial distribution functions showed that higher *ϵ*_b_ suppresses the population of configurations near the barrier region (***Figure 2—figure Supplement 1***C). This reduction in the statistical weight of barrier-region configurations can be interpreted as an entropy-related configurational restriction and thus disfavors phase separation. At the pair-potential level, the desolvation barrier modifies the equilibrium Boltzmann weight and thereby alters the integrated effective attraction, as quantified by the bead-level second virial coefficient *B*_2_. Specifically, increasing *ϵ*_b_ makes *B*_2_/*σ*^3^ larger (***Figure 2—figure Supplement 1***G), indicating a weaker integrated effective attraction and providing a thermodynamic basis for the lower 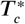. By contrast, deepening the solvent-separated well *ϵ*_ss_ elevates 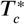 (Figure 2E), which is associated with the enhanced population of solvent-separated configurations and a smaller *B*_2_/*σ*^3^ (***Figure 2—figure Supplement 1***D, H). This solvent-separated minimum provides effective free-energy stabilization for water-mediated configurations and thereby promotes phase separation. These opposing effects suggest that the desolvation potential regulates macroscopic phase behavior by redistributing residue-pair configurations between direct-contact, barrier-region, and solventseparated states (lower panels of Figure 2D and E).

Because the critical temperature is an intrinsic, sequence-specific property of a given system, it is also important to examine how desolvation strength alters the overall shape of the phase diagram when the temperature is normalized by the corresponding 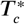. To do so, we rescaled the temperature axis by 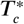, which effectively normalizes overall energetic strength and isolates desolvation-specific effects on the phase boundary. For the desolvation models with different barrier heights *ϵ*_b_, the normalized binodal curves nearly overlap (Figure 2F). By contrast, varying the solvent-separated well depth *ϵ*_ss_ results in distinct deviations in the density of the condensed phase (Figure 2G). Specifically, stronger water-mediated interactions lower equilibrium density by biasing the ensemble toward configurations with larger inter-residue separations (***Figure 2—figure Supplement 1***E, F). Thus, solvent-separated interactions directly regulate molecular packing and, by introducing solvent-separated configurations, alleviate the overcompaction typically encountered in coarse-grained models with implicit solvent. These results suggest that incorporating desolvation effects may enable the model to better capture the physicochemical structure of biological condensates.

### Conformational modulation by desolvation in dense and dilute phases

We next investigated how desolvation modulates the conformational distribution of IDPs across dilute and condensed phases. Previous studies have shown that IDP condensation can reorganize chain conformations by redistributing the balance between intra-chain and inter-chain interactions ***Wei et al. (2017); Hazra and Levy (2021); Tesei et al. (2021); von Bülow et al. (2025***). To quantify global chain dimensions, we calculated the radius of gyration (*R*_*g*_) as a primary structural metric. Consistent with this picture, the resulting *R*_*g*_ distributions show that protein chains adopt more compact conformations in the dilute phase, whereas chains in the condensed phase exhibit more extended conformations (Figure 3B and C). This conformational expansion upon condensation reflects a transition from intra-chain-dominated interactions in the dilute phase to inter-chain-dominated interactions in the dense phase ***Wohl et al. (2025***). Within the condensed phase, protein chains preferentially form inter-chain contacts that lower the free energy of the system, compensating for the entropic cost associated with reduced translational and conformational freedom.

**Figure 3.**
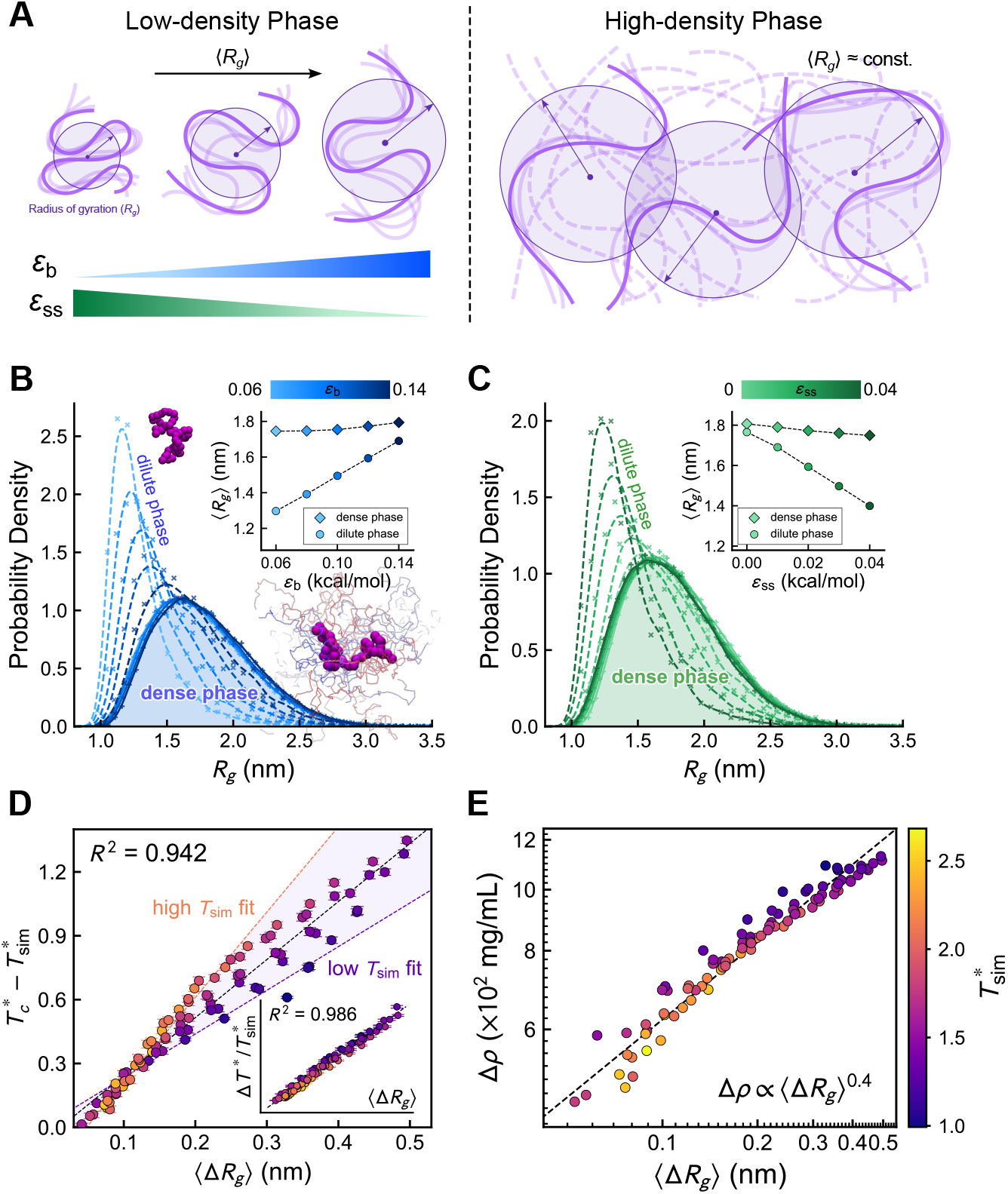
Effect of desolvation on protein conformations. (A) Schematic illustration of the conformational distributions of the protein in the high- and low-density phases under different desolvation parameters. (B–C) Distribution of *R*_*g*_ with different *ϵ*_b_ (B) and *ϵ*_ss_ (C) at 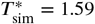. The solid and dashed lines represent the results for the condensed and dilute phases, respectively. The inset illustrates the mean value of *R*_*g*_ as a function of desolvation parameters. (D) Correlation between temperature difference 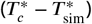 and the averaged *R*_*g*_ difference between two phases. The purple and yellow dashed lines represent linear fits to the data obtained at low- 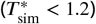 and high-simulation-temperature 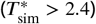 regimes, respectively. The inset displays the improved linearity obtained when the temperature difference is rescaled by the simulation temperature, 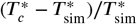. (E) Correlation between the density difference and *R*_*g*_ difference in the two phases with varying temperatures and desolvation parameters. The plot is shown on a log–log scale.

In addition to the difference between diluteand dense-phase conformations, varying the desolvation parameters further reveals a phase-dependent conformational response (Figure 3A–C). Increasing *ϵ*_b_ or decreasing *ϵ*_ss_ shifts the dilute-phase *R*_*g*_ distributions toward larger values, whereas the dense-phase *R*_*g*_ remains comparatively insensitive to these parameter changes. This contrast suggests that desolvation more strongly affects chain dimensions in the dilute phase, while conformations in the dense phase are less responsive, likely reflecting the constraints imposed by intermolecular packing in the condensate. As a result, the desolvation-dependent variation in 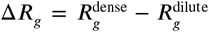 arises predominantly from the conformational changes of isolated chains in the dilute phase. Physically, Δ*R*_*g*_ captures the transition from intramolecular-dominated collapse in the dilute phase to intermolecular-mediated expansion in the dense phase, thus serving as a metric for the relative strength of inter-chain versus intra-chain interactions.

To further elucidate the thermodynamic implications of desolvation-modulated conformations, we analyzed the relationship between the conformational shift Δ*R*_*g*_ and the thermal distance to the critical point, defined as 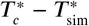. Notably, data from the simulated systems approximately follow a common trend, revealing a strong correlation between the magnitude of conformational change and the thermal distance to the phase transition point (*R*^2^ = 0.942, Figure 3D). This result suggests that the conformational response to phase separation is closely associated with how far the system resides thermally from the critical point.

We rationalize this observation using the Flory-Huggins theory ***Huggins (1942); Flory (1942***) as a bridge between the macroscopic thermal distance and the microscopic interaction strength. In this theoretical framework, the thermodynamic state is described by the interaction parameter *χ* = *ϵ*_eff_ /(*k*_*B*_*T*), where *ϵ*_eff_ represents the effective pairwise energy scale specific to each desolvation model ***Flory (1953***). At the critical point, *χ*(*T*_*c*_) = *χ*_*c*_, which gives *ϵ*_eff_ = *k*_*B*_*T*_*c*_*χ*_*c*_. Substituting this relation into the expression for *χ*(*T*_sim_) yields *χ*(*T*_sim_) = *χ*_*c*_*T*_*c*_/*T*_sim_. Subtracting *χ*_*c*_ from both sides gives

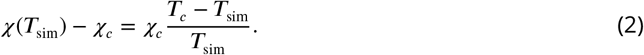

For systems with the same chain length, *χ*_*c*_ is a fixed constant. Thus, the deviation from the critical interaction parameter is directly related to the rescaled thermal distance (*T*_*c*_ − *T*_sim_)/*T*_sim_.

The thermodynamic driving force *χ*(*T*_sim_) − *χ*_*c*_ can then be related to the structural observable Δ*R*_*g*_. Since Δ*R*_*g*_ captures the structural transition from an intrachain-interaction-dominated state in the dilute phase to an interchain-interaction-dominated state in the dense phase, we assume, as a first-order approximation, that this conformational shift responds approximately linearly to the excess interaction strength, expressed as Δ*R*_*g*_ ∝ [*χ*(*T*_sim_) − *χ*_*c*_]. Combining these relations gives

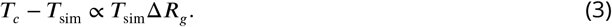

This relation explains the approximate linearity observed in Figure 3D, while also accounting for the systematic deviations caused by the temperature factor *T*_sim_, which is evident from the varying slopes at different simulation temperatures. To test this theoretical argument and the underlying linear-response assumption, we replotted the data against the rescaled thermal distance 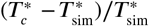. As shown in the inset of Figure 3D, this normalization yields a stronger linear relationship, supporting the above argument that Δ*R*_*g*_ scales with the excess interaction strength *χ* − *χ*_*c*_, and further suggesting that the conformational response is closely associated with the rescaled thermal distance from the critical point.

We further show the relationship between the phase density difference (Δ*ρ* = *ρ*_dense_ − *ρ*_dilute_) and the conformational difference (Δ*R*_*g*_) between coexisting phases in Figure 3E. Based on the empirical correlation between Δ*R*_*g*_ and the thermal distance from the critical point, one can relate the density contrast Δ*ρ* to Δ*R*_*g*_ through the critical-scaling relation by Δ*ρ* = *A*(*T*_*c*_ − *T*)^*β*^ (Equation (7)), where *β* is the critical exponent. Although the complete relation in Equation (3) contains an additional *T*_sim_ factor, the unscaled quantities remain strongly correlated over the simulated range. We therefore use *T*_*c*_ −*T*_sim_ ∝ Δ*R*_*g*_ as an empirical scaling approximation. Using this approximation in the critical-scaling relation yields Δ*ρ* ∝ (Δ*R*_*g*_)^*β*^ . On logarithmic scales, a clear power-law dependence emerges (Figure 3E), with 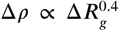, which is consistent with our theoretical expectation. The fitted exponent (≈ 0.4) is in reasonable agreement with the theoretical critical exponent *β* = 0.325 based on the 3D Ising model ***Rowlinson and Widom (2013***). Collectively, these results reveal a fundamental coupling between microscopic conformational reorganization and macroscopic packing density during biomolecular condensation, highlighting conformational reorganization as a structural signature of the phase transition.

### Dynamic consequences of desolvation in phase separation

Beyond governing thermodynamic phase boundaries and conformational distribution, the desolvation potential fundamentally reshapes the dynamical landscape of IDP condensates. The interplay between inter-chain interactions and solvent exclusion not only dictates the stability of the dense phase but also modulates the transport properties within the condensate and the macroscopic kinetics of phase separation. To probe chain mobility within the dense phase, we employed a strategy analogous to fluorescence recovery after photobleaching (FRAP) experiments following prior molecular dynamics simulation work ***Yamada and Takada (2023***), tracking the spatiotemporal evolution of specific chains initially localized at the slab center (Figure 4A and B). The systems were compared at the same simulation temperature 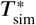. For the standard HPS model without desolvation, the tagged chains achieve a uniform distribution that matches the equilibrium density profile within 50 ns (Figure 4B), consistent with the liquid-like nature of the condensate. For the representative desolvation model, the spatial distribution of tracked chains remains biased toward the initial center position after 50 ns and approaches equilibrium on a prolonged timescale (∼ 150 ns). These contrasting relaxation profiles demonstrate that desolvation alters chain mobility within the condensate.

**Figure 4.**
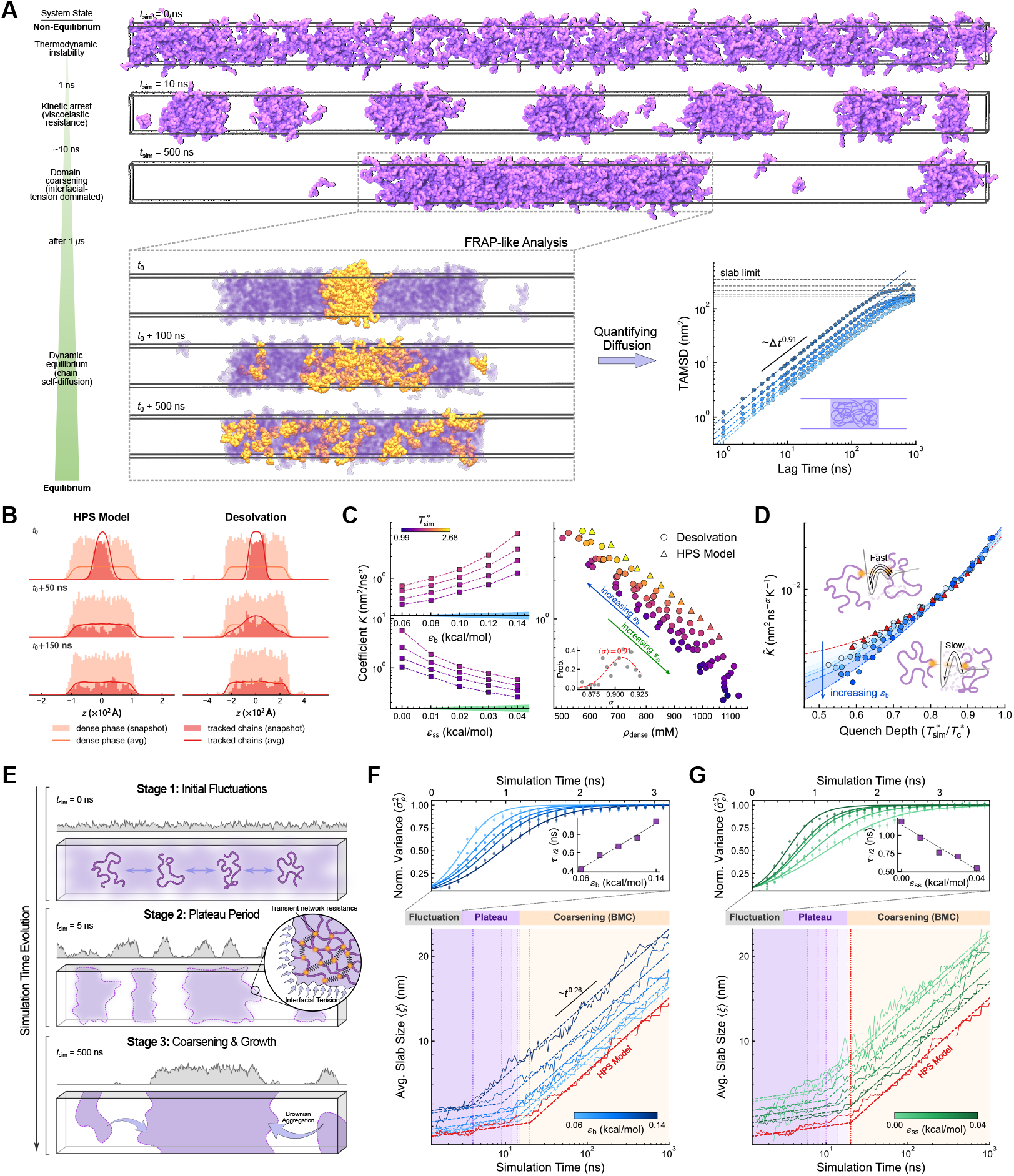
Desolvation-mediated modulation of diffusion and coarsening dynamics. (A) Snapshots of phase-separation dynamics after a temperature quench and the subsequent FRAP-like analysis of chain self-diffusion in an equilibrated slab. Upper snapshots show the simulation box along the *z*-axis at *t*_sim_ = 0, 10, and 500 ns. The vertical schematic summarizes the dynamical progression described in the main text, from the post-quench spinodal instability to kinetic arrest, domain coarsening, and dynamic equilibrium. Lower panels show enlarged slab views for tracking chains initially located near the slab center, with the TAMSD plot quantifying their mobility. (B) Density profiles along the *z*-axis for the entire system (orange) and for the highlighted chains (red) at different time lags, with and without desolvation. Shaded histograms show instantaneous snapshots, and solid curves represent normalized time averages over 1 *μ*s. (C) Diffusion coefficients as a function of the desolvation strength (*ϵ*_b_, *ϵ*_ss_) and dense-phase density (*ρ*_dense_) at different temperatures. (D) Reduced diffusion coefficient 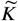 versus quench depth for varying *ϵ*_b_. The schematic insets summarize the reduced chain mobility observed for higher desolvation barriers at comparable quench depths. (E) Schematic illustration of the three-stage phase-separation mechanism, including early density-fluctuation growth, a transiently arrested plateau, and late-stage coarsening. In the zoom-in view, yellow beads denote residues involved in transient inter-chain contacts, and black springs denote schematic network connections formed by these contacts. (F, G) Kinetics of density fluctuations and domain growth under varying *ϵ*_b_ (F) and *ϵ*_ss_ (G). Top panels show the time evolution of the normalized density variance 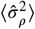, with the inset showing the characteristic time *τ*_1/2_. Bottom panels show the growth of the average slab size ⟨*ξ*⟩ on a logarithmic scale. The colored timeline highlights the initial fluctuation, plateau, and late-stage coarsening regimes.

To more quantitatively assess the influence of desolvation on chain mobility, we computed the time-averaged mean squared displacement (TAMSD) ***Metzler et al. (2014***) for chains within the slab. The chain motion exhibits sub-diffusive scaling, TAMSD = *K* ⋅ Δ*t*^*α*^, with a characteristic exponent *α* ≈ 0.91 (Figure 4A, bottom right panel). This deviation from normal diffusion (*α* = 1) likely reflects the combined effects of molecular crowding, multivalent dynamic interaction networks, and finite-size confinement within the dense phase. At extended lag times, the TAMSD curves naturally plateau as they approach the geometric limit imposed by the slab thickness. Notably, the observed exponent is comparable to simulation results reported for protein diffusion in threedimensional condensates (*α* ≈ 0.85) ***Watanabe et al. (2025***), consistently highlighting the restricted nature of chain dynamics within the dense phase. Given the relatively consistent values of *α* across different interaction parameters, we use *K* as a quantitative measure of the chain diffusion rate.

The diffusion prefactor *K* exhibits a clear dependence on the interaction parameters (Figure 4C). At a fixed temperature, *K* increases with the desolvation barrier *ϵ*_b_ and decreases with the solvent-separated minimum depth *ϵ*_ss_ (Figure 4C, left panels). These opposing trends can be largely understood from the distinct effects of the two desolvation parameters on condensate packing. As illustrated in Figure 4C (right panel), *K* is predominantly correlated with the bulk density (*ρ*_dense_), with further modulation by temperature. As expected, higher-temperature systems generally exhibit slightly larger *K* values at comparable densities. Consistent with the density trends established in Figure 2D and E, a higher *ϵ*_ss_ promotes tighter packing and increases the effective friction, whereas a higher *ϵ*_b_ prevents close contacts and creates greater free volume that facilitates chain motion. These results indicate that, at a fixed absolute temperature, changes in dense-phase packing serve as the primary determinant of chain mobility.

However, this density-dominated view is incomplete because it convolutes intrinsic energy-landscape features with thermodynamic state shifts. Since variations in *ϵ*_b_ and *ϵ*_ss_ significantly alter the critical temperature, comparisons made at a constant simulation temperature inherently involve systems positioned at different thermodynamic quench depths, defined as the normalized temperature deviation from the critical point. This highlights the distinct thermodynamic driving forces for different systems. To reduce this thermodynamic-state dependence, we performed a more carefully controlled comparison. First, we scaled the simulation temperature by the critical temperature of each system and used 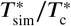 as the control parameter for quench depth, with a lower value corresponding to a deeper quench. This provides more comparable thermodynamic conditions across the different interaction parameters. Second, to account for the leading linear temperature dependence associated with Langevin dynamics, we defined a reduced diffusion prefactor 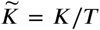 as detailed in the Methods section. Under these controlled conditions, a clearer parameter-dependent kinetic trend emerges (Figure 4D). At the same quench depth, a higher desolvation barrier *ϵ*_b_ (darker blue points) lowers 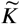, consistent with a rougher energy landscape that hinders the configurational rearrangements required for chain diffusion. This kinetic divergence becomes more pronounced at deeper quenches but largely vanishes near the critical point, indicating that the kinetic influence of the desolvation barrier depends on the thermodynamic state. Notably, the standard HPS model exhibits much faster temperature-normalized mobility (Figure 4D, red triangles), suggesting that omitting desolvation energetics may lead to an overestimation of chain mobility due to reduced roughness of the effective energy landscape.

Together, the fixed-temperature and renormalized analyses in Figure 4C and D reveal two distinct and opposing contributions of desolvation to condensate dynamics. At fixed temperature, increasing *ϵ*_b_ loosens dense-phase packing and thereby increases the measured diffusion coefficient, whereas at matched thermodynamic quench depth, the same parameter change suppresses chain mobility by roughening the microscopic energy landscape and slowing local rearrangements. Condensate dynamics therefore emerge from a balance between density-regulated mobility and energy-landscape-regulated mobility, with macroscopic packing determining the dominant trend and microscopic barrier roughness imposing an additional kinetic modulation. This interplay highlights how desolvation reshapes condensate dynamics across multiple physical scales.

### Desolvation-modulated kinetic arrest of coarsening dynamics

Elucidating the molecular events during the early stage of phase separation is challenging experimentally. We therefore examined droplet-growth dynamics at the early stage of phase separation by performing molecular simulations with desolvation-aware effective potentials. Initially, the system was equilibrated at a supercritical temperature (*T* ^∗^ = 2.98) for 10 ns to ensure a homogeneous distribution. The temperature was then quenched to *T* ^∗^ = 1.79, located deep within the two-phase coexisting region, to initiate spontaneous phase separation (Figure 4A). Here, *t*_sim_ = 0 ns denotes the first snapshot immediately after the temperature quench, corresponding to a homogeneous but thermodynamically unstable non-equilibrium state. During the subsequent time evolution, the initially uniform distribution rapidly develops interconnected density fluctuations within 1 ns to 2 ns, which then break up into discrete high-density domains along the *z*-axis. Over time, these domains gradually coalesce into larger slabs, exhibiting the characteristic morphological evolution of spinodal decomposition ***Cahn and Hilliard (1958***).

To quantify the phase separation kinetics, we adopted a two-stage metric strategy. In the early stage, where interfaces are diffusive, we monitored the normalized density variance (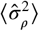, see Supplementary Information) derived from the instantaneous density profile *ρ*(*z*) to track the growth rate of density fluctuations. As clear interfaces emerged, we switched to tracking the average slab size (⟨*ξ*⟩), defining dense domains as continuous regions where *ρ*(*z*) exceeds a threshold value (here the average system density was used as the threshold). To ensure statistical robustness, all kinetic metrics were averaged over six independent slab simulation replicas. The resulting kinetic profiles (Figure 4F and G) resolve a three-stage dynamical pathway. In the first stage, immediately after the temperature quench, the system undergoes a rapid exponential growth in density variance (top panels). This behavior arises from the growth of initial density fluctuations in a thermodynamically unstable state and is characteristic of spinodal decomposition ***Cahn (1961***). Stronger effective attraction, corresponding to higher *ϵ*_ss_ or lower *ϵ*_b_, accelerates these initial fluctuations, as quantified by the monotonic decrease in the characteristic time scale *τ*_1/2_, defined as the time required to reach 50% of the maximum normalized variance.

Intriguingly, after dense domains have formed, the system enters an intermediate plateau regime in which domain growth is temporarily arrested before late-stage coarsening begins. This kinetic arrest is evident from the slab-size evolution in Figure 4F and G, where ⟨*ξ*⟩ remains nearly unchanged for a distinct period, and is visually corroborated by the density profiles at 5 ns in Figure 4E, where distinct domains are observed to form but fail to fuse immediately. This behavior is consistent with the characteristics of viscoelastic phase separation ***Tanaka (2000***). In this regime, interfacial tension favors domain fusion, whereas transient inter-chain connectivity within the dense domains generates viscoelastic resistance to the shape deformation required for coalescence, thereby delaying domain fusion. Similar viscoelastic effects have recently been quantified in molecular simulations of biomolecular condensates using rheological analyses of time-dependent material properties ***Tejedor et al. (2023***). Molecular simulations of condensate aging have further highlighted the roles of chain flexibility, sticker lifetime, and desolvation-associated rigidification in promoting more solid-like states ***Biswas and Potoyan (2024***). In the zoom-in schematic in Figure 4E, this transient network is represented by connections between residues involved in inter-chain contacts, illustrating how multivalent interactions can resist domain deformation during the plateau regime. Quantitative analysis in Figure 4F and G further reveals that the duration of this plateau regime depends strongly on the desolvation parameters. Systems with higher *ϵ*_ss_ or lower *ϵ*_b_ exhibit a longer plateau, indicating that stronger inter-chain attractions reinforce the transient network and prolong kinetic arrest. This trend corroborates our viscoelastic interpretation, as enhanced intermolecular connectivity increases the resistance to domain deformation and fusion. This kinetic trap is particularly pronounced in the standard HPS model (red trajectories in Figure 4F and G), consistent with its relatively stronger pairwise attractions.

At later times (*t* > 10 ns), the system leaves the plateau regime and enters a coarsening-and-growth stage, where domain fusion proceeds through interfacial-tension-driven relaxation and Brownian aggregation. The average slab size follows a power-law scaling ⟨*ξ*⟩ ∼ *t*^*α*^, with *α* ≈ 0.26 in our simulations. This exponent is slightly lower than the asymptotic prediction of Brownian motion coalescence (BMC), *α* = 1/3 ***Majumdar et al. (1994***), likely because the viscoelastic nature of the polymer-rich phase and the sub-diffusive motion of individual chains (TAMSD ∼ Δ*t*^0.91^) constrain macroscopic domain diffusion ***Doi et al. (1988***).

Collectively, these analyses reveal a desolvation-modulated kinetic pathway that links early density fluctuations, transient domain arrest, and late-stage coarsening into a unified dynamical picture, as summarized by the vertical schematic in Figure 4A and the mechanistic illustration in Figure 4E. This picture shows that desolvation does more than slow down local chain rearrangements through an added barrier. It also regulates the balance between fluctuation growth, transient arrest, and domain coarsening, thereby shaping the evolution of phase-separated domains. The sensitivity of kinetic arrest and coarsening dynamics to desolvation parameters underscores the importance of incorporating desolvation features into coarse-grained potentials for more physically plausible molecular simulations of LLPS, especially when connecting microscopic interaction lifetimes to emergent viscoelastic or aging-like material behavior. Beyond modulating LLPS thermodynamics and conformational ensembles as discussed above, desolvation therefore acts as a critical regulator of phase-separation dynamics.

### Parameterization of desolvation model for IDPs

The results from the homopolymer model discussed above underscore the critical role of desolvation in regulating both the thermodynamics and dynamics of LLPS. Consequently, incorporating water-mediated interactions into residue-level coarse-grained models is essential for capturing the physics of protein phase separation more realistically. To this end, we developed a transferable parameterization strategy that combines information derived from all-atom simulations with calibration against experimental observables of IDPs.

Directly determining residue-specific desolvation parameters, specifically the barrier height *ϵ*_b_ and the solvent-separated minimum depth *ϵ*_ss_, remains challenging. To bridge this gap, we leveraged all-atom molecular dynamics simulations of amino acid analogues. Using the global interaction energy scale *ϵ* of the underlying HPS-type potential as a reference, we expressed the desolvation amplitudes as *ϵ*_b_ = *α*_b_*ϵ* and *ϵ*_ss_ = *α*_ss_*ϵ*. By fitting the PMF profiles obtained from all-atom simulations (***Figure 1—figure Supplement 1***), we obtained coefficients of comparable magnitude across the limited set of analogue pairs examined. For simplicity and transferability, we therefore adopted the averaged values *α*_b_ = 0.33 and *α*_ss_ = 0.06 as a global baseline for our coarse-grained model. This uniform parameterization captures the generic desolvation features of the PMFs but does not resolve residue-pair-specific variations in desolvation energetics.

To evaluate the effectiveness of these physics-derived parameters, we first integrated them into the framework of the standard HPS model. While the HPS model performs remarkably well in capturing sequence-dependent properties of LLPS, previous studies have noted that parameters optimized solely against single-chain *R*_*g*_ may have limited transferability to the condensed phase, resulting in an overestimation of phase-separation propensity, as reflected by higher critical temperatures (*T*_*c*_) ***Dignon et al. (2018b***). We assessed the model performance on a dataset of 47 IDPs taken from Tesei et al. ***Tesei et al. (2021***), which was subsequently used to calibrate the CALVADOS2desolvation parameters. The deviation of simulated single-chain *R*_*g*_ from experimental SAXS data was quantified by 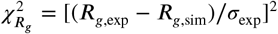, where *σ*_exp_ represents the experimental uncertainty. When the desolvation parameters derived from all-atom simulations (*α*_b_ = 0.33, *α*_ss_ = 0.06) were added to the HPS model without modifying its original *ϵ*, we observed a substantial improvement in conformational accuracy. The average discrepancy 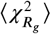 decreased drastically from 1,082.27 (original HPS) to 85.88 (HPS-desolvation) (Figure 5B). This improvement indicates that introducing the desolvation terms effectively mitigates the over-compaction tendency of the original HPS model, likely because of competition between direct and water-separated contacts.

**Figure 5.**
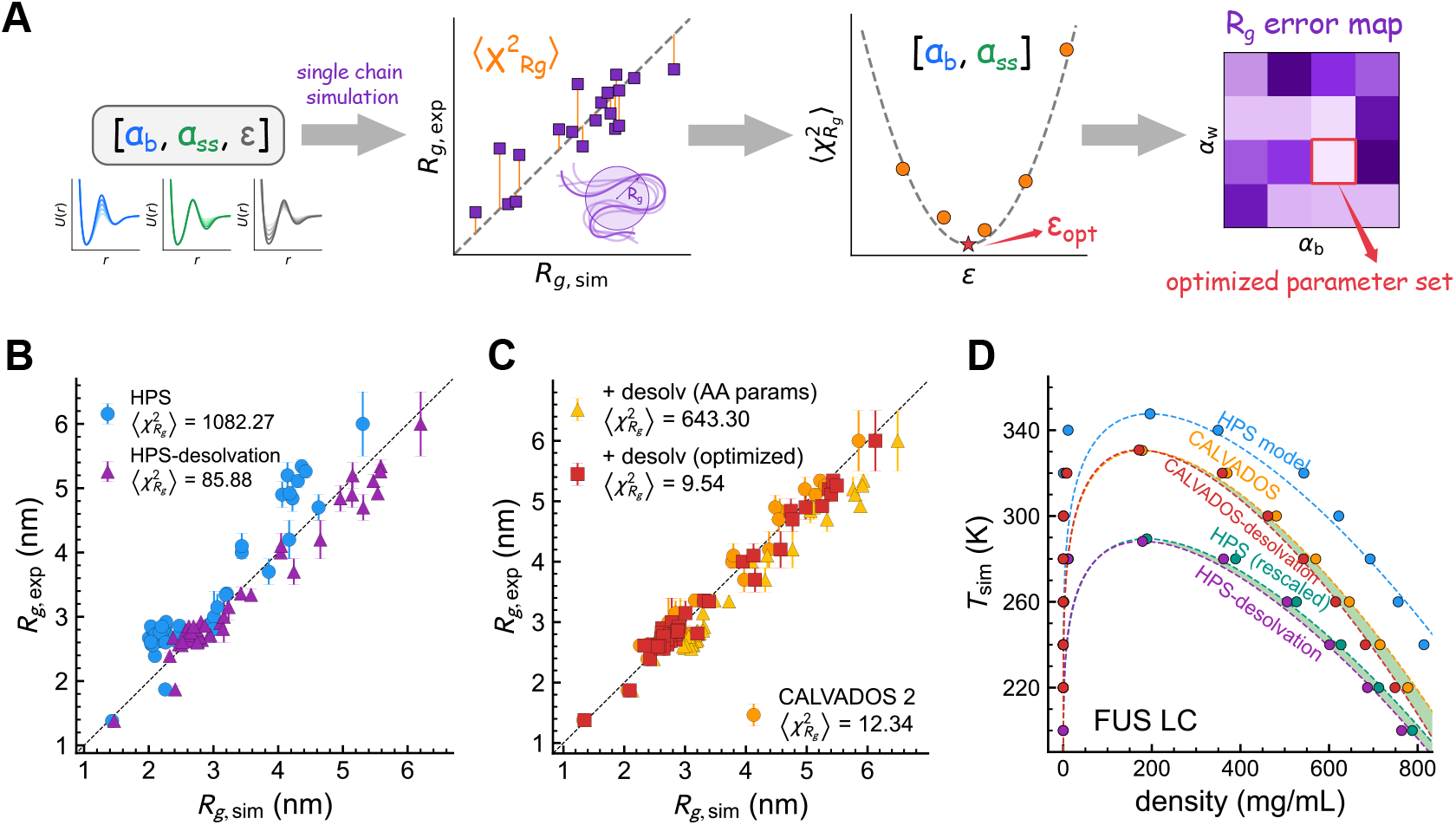
Parameterization of desolvation terms for the HPS and CALVADOS2 models based on IDPs. (A) Schematic workflow of the desolvation parameterization. (B) Correlation between experimental *R*_*g*_ and simulation *R*_*g*_ for the original HPS model (blue) and the revised HPS model with default desolvation scales (*α*_b_ = 0.33 and *α*_ss_ = 0.06) (purple). The 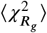 for different models are also shown. (C) Correlation between experimental *R*_*g*_ and simulation *R*_*g*_ for the original CALVADOS2 model (orange), the revised CALVADOS2 model with default desolvation (yellow), and the revised CALVADOS2 model with optimized desolvation (*α*_b_ = 0.3, *α*_ss_ = 0.03 and *ϵ* = 0.262 kcal/mol, red). (D) Coexistence curves of FUS LC simulated with the original HPS model (blue), the revised HPS model with default desolvation scales (purple), the energy-rescaled HPS model (green), the default CALVADOS2 model (orange), and the revised CALVADOS2 model with optimized desolvation scales (red). The green shaded regions highlight the deviations of the desolvation models relative to the original frameworks.

We also incorporated the desolvation terms into CALVADOS2 ***Tesei and Lindorff-Larsen (2022***), a force field whose hydrophobicity parameters (*λ*) were reparameterized using Bayesian optimization to reproduce experimental *R*_*g*_ and PRE data. Directly adding desolvation terms to CALVADOS2 led to substantial chain over-expansion 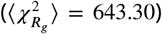, reflecting the fact that CALVADOS2 is already finely tuned to reproduce experimental data through a top-down optimization procedure and the inclusion of additional desolvation terms is incompatible with its parameterization. To address this, we implemented a systematic optimization strategy in which the desolvation barrier height (*ϵ*_b_), solvent-separated minimum depth (*ϵ*_ss_), and the overall energy factor *ϵ* were scanned over broad parameter ranges to identify the parameter set that minimizes deviations between simulated and experimental *R*_*g*_ values. The full optimization workflow is summarized in the flowchart in Figure 5A, and the resulting error landscapes are shown in ***Figure 5—figure Supplement 1*** and ***Figure 5 figure Supplement 2*** . The optimized parameter set (Table 1) reduced 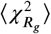 to 9.54 on the calibration dataset (Figure 5C), yielding agreement with experimental *R*_*g*_ values comparable to that of the original CALVADOS2 model.

**Table 1.** Global desolvation coefficients *α*_b_ and *α*_ss_ used in the HPS and CALVADOS2 frameworks. The HPS baseline coefficients were obtained by averaging values fitted to the all-atom analogue PMFs, whereas the CALVADOS2 coefficients and global energy scale *ϵ* were selected through optimization against experimental *R*_*g*_ data.

| Parameters | HPS model | CALVADOS2 model |
| --- | --- | --- |
| $\alpha_b$ | 0.33 | 0.30 |
| $\alpha_{ss}$ | 0.06 | 0.03 |
| $\epsilon$ (kcal/mol) | 0.20 | 0.262 |

After parameterization of the energy scale, we further evaluated the calibrated model by probing the phase behavior of the FUS low-complexity domain (FUS LC) as a test case. Experimental observations indicate that the critical temperature is around 25 ^°^C ***Burke et al. (2015***). While the original HPS model yields a critical temperature of *T*_*c*_ ∼ 340 K, incorporating the desolvation terms into the HPS framework (HPS-desolvation) lowers the model *T*_*c*_ to ∼ 290 K (Figure 5D), shifting it toward the experimentally observed room-temperature regime. It is worth noting that the residue-level coarse-grained models used here employ temperature-independent effective interaction parameters. As a result, the temperature values reported here cannot be interpreted as quantitatively equivalent to experimental temperatures, particularly when they deviate substantially from room-temperature conditions. Beyond this reduction in the excessive phase-separation propensity, we observed that the desolvation model mitigates the overestimation of condensate density common in implicit solvent coarse-grained simulations ***Regy et al. (2021); Benayad et al. (2020***). In the CALVADOS2 framework, the inclusion of desolvation reduces the dense phase density while maintaining an overall phase boundary comparable to the original model (Figure 5D). Similarly, for the HPS model, when we rescaled the interaction strength *ϵ* to match the single-chain *R*_*g*_ and thereby align the phase boundary with the HPS-desolvation system (***Figure 5—figure Supplement 3***), we observed the same behavior. A similar trend was observed in the simulations of the LAF-1 RGG domain, where CALVADOS2-desolvation preserved a comparable coexistence curve while modestly reducing the dense-phase density relative to the original CALVADOS2 model (***Figure 5—figure Supplement 4***). Quantitatively, across both frameworks, the desolvation-modulated models consistently yield a packing density approximately 5% lower than the original models. This distinction underscores a limitation of standard Lennard-Jones potentials where a single parameter *ϵ* simultaneously controls chain dimension, overall phase-separation propensity, and condensate density. In contrast, our model introduces tunable parameters *ϵ*_b_ and *ϵ*_ss_ that provide an additional physical handle for modulating dense-phase packing without simply rescaling the overall interaction strength. The observed density reduction can be attributed to an increased population of solvent-separated configurations relative to direct contacts. This effect provides an effective representation of hydration-associated packing in protein condensates, thereby offering a more physically realistic description of IDP phase separation.

## Discussion and Conclusion

Coarse-grained molecular simulations have become indispensable for elucidating the molecular principles underlying LLPS, complementing both theoretical frameworks and experimental observations. While residue-level implicit-solvent models have achieved remarkable success in capturing sequence-dependent phase behavior, they usually do not separately account for desolvation effects associated with water-mediated residue-residue association. As a result, they often face a trade-off when simultaneously describing single-chain dimensions, phase-separation propensity, and dense-phase packing. Specifically, interaction parameters calibrated to match single-chain dimensions or phase-separation propensity frequently result in an overestimation of the condensed phase density. This limitation suggests that standard Lennard-Jones potentials, which couple overall phase-separation propensity and dense-phase packing through a single energetic parameter, may lack the physical dimensionality required to fully describe the thermodynamics of hydrated protein condensates. The present results therefore suggest that desolvation is not merely a microscopic correction to residue-residue contacts, but an additional physically motivated component in coarse-grained interaction models that can modulate LLPS thermodynamics, dense-phase packing, conformational reorganization, and condensate dynamics in partially separable ways.

To address this challenge, we developed a desolvation-aware implicit-solvent CG framework that incorporates desolvation barrier and solvent-separated terms into the pairwise potential. Application of this model to homopolymer systems demonstrates that desolvation substantially reshapes phase behavior. Incorporating desolvation modulates the critical temperature and alters the phase diagram by regulating intermolecular packing within the dense phase. Moreover, the model reveals a tight coupling between desolvation-controlled chain compaction during the formation of the condensate phase and the thermal distance from criticality (temperature gap), providing a theoretical basis for linking single-chain conformational changes (Δ*R*_*g*_) to macroscopic phase behavior. Dynamic analysis further captures the three characteristic stages of early LLPS, including spinodal decomposition, kinetic arrest, and coarsening. Crucially, it reveals how the desolvation effect regulates the kinetics of these stages by modulating the effective inter-chain interactions and the roughness of the energy landscape. These results underscore the importance of accounting for desolvation energetics in governing both the thermodynamics and kinetics of LLPS. These findings may also provide a useful physical basis for future studies of pressure-dependent condensate behavior, as pressure-induced changes in hydration, solvent-separated states, and desolvation barriers have been proposed to contribute to pressure-modulated and reentrant LLPS ***Dias and Chan (2014); Cinar et al. (2019***, 2018).

Motivated by these insights, we further developed a transferable parameterization strategy for residue-level models. The resulting desolvation-aware CG model improves the accuracy of IDP conformational ensembles and LLPS thermodynamics while preserving the computational efficiency inherent to CG methodologies. By retaining key water-mediated features while preserving the computational efficiency of implicit-solvent representations, this framework provides a mechanistic means to decouple overall phase-separation propensity from condensed-phase packing. As illustrated in Figure 5D, incorporating desolvation into CALVADOS2 reduces *ρ*_dense_ while preserving an overall phase boundary comparable to that of the original model. This decoupling effectively mitigates the overcompaction of the dense phase commonly observed in implicit solvent simulations with CG models, thereby making the simulations qualitatively more consistent with the physical nature of biological condensates.

Despite these advances, several limitations remain in the present implementation. First, the current model introduces desolvation terms on top of residue-level CG frameworks with simplified interaction parameterizations. In particular, HPS-type models use a 20-parameter hydropathy-scale representation, which is useful for capturing generic IDP phase behavior but is not flexible enough to resolve residue-pair-specific chemical effects, such as the distinct interaction patterns of arginine and lysine residues ***Das et al. (2020***). More general 210-parameter pairwise interaction schemes, such as KH-type and related residue-pair-specific models, provide greater flexibility for encoding amino-acid-pair preferences and capturing sequence-specific interaction heterogeneity ***Dignon et al. (2018b); Joseph et al. (2021); Wessén et al. (2022***). Second, the present desolvation model employs a single set of averaged parameters (*α*_b_, *α*_ss_) for all residue pairs. While this simplification is effective for isolating the generic physical consequences of desolvation, it has limitations in describing the pair-specific variations in the desolvation barrier and the solvent-separated minimum. Explicit-solvent PMF analyses have shown that desolvation barrier heights and solvent-separated minima can differ substantially among residue pairs and may also exhibit temperature dependence ***Cinar et al. (2019); Debiec et al. (2014); MacCallum et al. (2007***). In addition, effective interactions themselves can be temperature dependent, as demonstrated in analytical theories and coarse-grained simulations of biomolecular condensates ***Lin et al. (2017); Dignon et al. (2019); Chakravarti and Joseph (2025***). Future extensions of the model could therefore incorporate residue-specific and temperature-dependent desolvation parameters derived from bottom-up parameterization or expanded experimental datasets, thereby enhancing predictive accuracy for sequence-dependent LLPS. An additional direction would be to combine these potentials with simulation-based rheological analyses to quantify how desolvation reshapes condensate viscoelasticity, aging-like maturation, and long-time material relaxation ***Tejedor et al. (2023); Biswas and Potoyan (2024***).

In summary, this work provides a physically interpretable route for incorporating solvent-mediated interaction features into implicit-solvent CG models, providing a basis for future residue-pair-specific and condition-dependent models of biomolecular condensates. By enabling an efficient representation of key desolvation-related features, the model provides a robust framework for simulating biomolecular condensates with improved thermodynamic, structural, and kinetic fidelity. Future developments, such as residue-pair-specific desolvation energetics and extensions to multicomponent systems involving nucleic acids, hold strong potential for deepening our mechanistic understanding of the complex phase behaviors that organize cellular biochemistry.

## Methods

### All-atom MD simulations

The all-atom MD simulations were performed using GROMACS. Ten amino acid analog molecules were solvated in a cubic water box under periodic boundary conditions. The OPLS-AA force field was used to model the solute molecules, and the TIP4P water model was employed to represent solvent. Na^+^ and Cl^−^ ions were added to neutralize the system, yielding a total system size of 1,010 atoms. After an initial energy minimization of 5 × 10^4^ steps, the system was equilibrated in the NVT ensemble at 298 K for 100 ps, followed by an NPT equilibration at 298 K and 1 atm for 1 ns. Temperature and pressure were controlled using the Nosé–Hoover thermostat and Parrinello– Rahman barostat ***Parrinello and Rahman (1981***), respectively. A time step of 2 fs was used for all MD steps. The simulation details are provided in Table 2. Subsequently, a production simulation of 100 ns was performed under the same NPT conditions for structural and thermodynamic analysis. The intermolecular separation *r* between two amino acid analogues was defined as the distance between their C atoms. The potential of mean force (PMF) was computed as PMF(*r*) = −*k*_B_*T* ln *P* (*r*), where *P* (*r*) is the radial probability density obtained from the production trajectory.

**Table 2.**
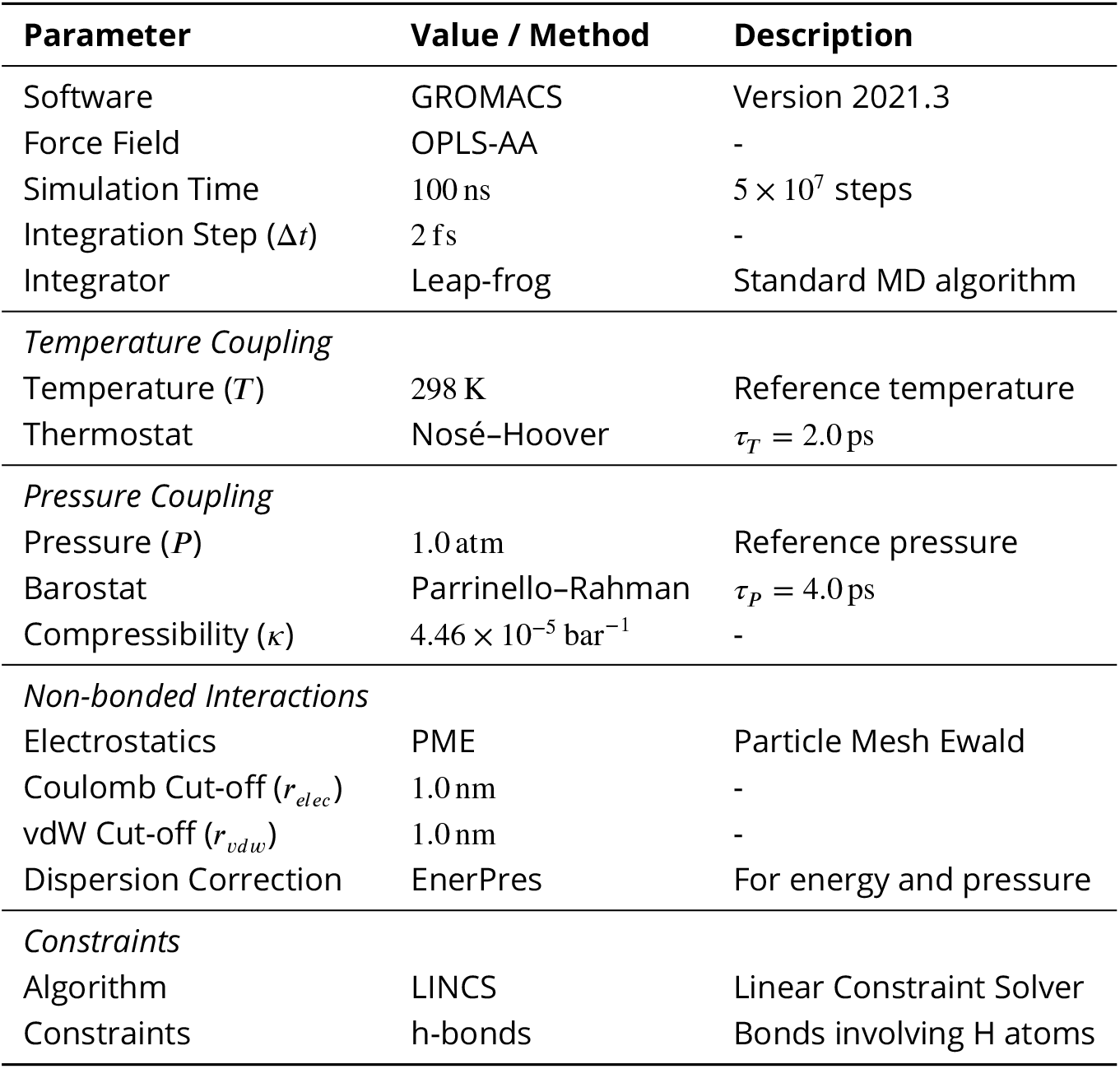
Summary of the production run parameters for all-atom molecular dynamics simulations.

| Parameter | Value / Method | Description |
| --- | --- | --- |
| Software | GROMACS | Version 2021.3 |
| Force Field | OPLS-AA | - |
| Simulation Time | 100 ns | $5 \times 10^7$ steps |
| Integration Step ($\Delta t$) | 2 fs | - |
| Integrator | Leap-frog | Standard MD algorithm |
| *Temperature Coupling* |  |  |
| Temperature ($T$) | 298 K | Reference temperature |
| Thermostat | Nosé-Hoover | $\tau_T = 2.0$ ps |
| *Pressure Coupling* |  |  |
| Pressure ($P$) | 1.0 atm | Reference pressure |
| Barostat | Parrinello-Rahman | $\tau_P = 4.0$ ps |
| Compressibility ($\kappa$) | $4.46 \times 10^{-5}$ bar$^{-1}$ | - |
| *Non-bonded Interactions* |  |  |
| Electrostatics | PME | Particle Mesh Ewald |
| Coulomb Cut-off ($r_{elec}$) | 1.0 nm | - |
| vdW Cut-off ($r_{vdw}$) | 1.0 nm | - |
| Dispersion Correction | EnerPres | For energy and pressure |
| *Constraints* |  |  |
| Algorithm | LINCS | Linear Constraint Solver |
| Constraints | h-bonds | Bonds involving H atoms |

### Coarse-grained simulations

We utilized the slab method ***Dignon et al. (2018b***) to extract the densities of the coexisting dilute and dense phases. In these simulations, 125 protein chains were placed in a rectangular box of dimensions 15 nm × 15 nm × 280 nm. The density profile along the *z*-axis was computed to identify the condensed and dilute regions, from which thermodynamic quantities such as the critical temperature were subsequently determined. Simulations were performed over a range of different temperatures. To incorporate the desolvation terms into the pairwise interaction potentials of the HPS model, we used the azplugins package (https://github.com/mphowardlab/azplugins) together with HOOMD-blue v2.9.7 ***Anderson et al. (2020***). We also implemented the desolvation terms for both the HPS and CALVADOS models based on the OpenMM open-source toolkit ***Eastman et al. (2023***). GPU-accelerated simulations were performed on NVIDIA GeForce RTX 4090D and Tesla V100 GPUs, and CPU-based simulations were carried out on Kunpeng 920B. All simulations used a time step of 0.01 ps. Each trajectory was first equilibrated for 3 × 10^7^ steps, followed by a production run of 1 × 10^8^ steps. Additional details of the simulation framework are provided in Table 3.

**Table 3.** Summary of the coarse-grained molecular dynamics simulation parameters used in the slab simulations.

| Parameter | Value / Method | Description |
| --- | --- | --- |
| *General Settings* |  |  |
| Software | HOOMD-blue<br>OpenMM | v2.9.7 with azplugins<br>v8.2.0 |
| Integrator | Langevin | Stochastic dynamics |
| Time Step ($\Delta t$) | 0.01 ps | - |
| Total Production Time | 1 μs | 10<sup>8</sup> steps |
| Temperature ($T$) | Variable | See main text |
| Langevin Damping Rate ($\gamma_i/m_i$) | 0.01 ps<sup>-1</sup> | Mass-dependent |
| *System Geometry* |  |  |
| Box Dimensions ($x, y$) | 15.0 nm | Periodic boundaries |
| Box Dimension ($z$) | 280.0 nm | Slab configuration |
| *Interactions* |  |  |
| Bond Potential | Harmonic | $k = 8368 \text{ kJ} \cdot \text{mol}^{-1} \cdot \text{nm}^{-2}$, $r_0 = 0.38 \text{ nm}$ |
| Non-bonded Potential | Ashbaugh-Hatch | HPS model (via azplugins) |
| Non-bonded Cut-off | 2.4 nm | - |
| Electrostatics | Yukawa | Screened Coulomb potential |
| Screening Constant ($\kappa$) | 1.0 nm<sup>-1</sup> | Inverse Debye length |
| Coulomb Cut-off | 3.5 nm | - |
| Neighbor List | Cell List | Exclusions: 1-2, body |

### Hydrophobicity Scale Model

Our desolvation model was developed based on the hydrophobicity scale (HPS) coarse-grained model by Dignon et al. ***Dignon et al. (2018b***), in which each residue is represented by a single bead. The potential energy comprised a bonded term, an electrostatic term and a short-range pairwise term. The bonded interaction term is given by a harmonic potential *V* ^*b*^ = *k*_*b*_(*r*_*i*,*i*+1_ − *r*^0^)^2^ with the spring constant *k*_*b*_ = 10 kcal/mol/Å^2^ and the bonded length *r*^0^ = 3.8 Å. *r* _*i*,*i*+1_ represents the distance between the consecutive residues *i* and *i* + 1 along the protein chain. The electrostatic interaction term is modeled by a Coulombic potential with Debye-Hückel ***Debye and Hückel (1923***) electrostatic screening caused by salt ion concentration. The function is given by

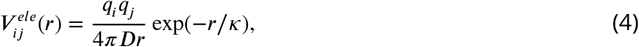

where *κ* is the screening length and *D* = 80, which is the relative dielectric constant of water. For the short-range pairwise interaction, the HPS model uses a 6-12 Lennard-Jones potential with the interaction strength given by hydrophobicity scale ***Kapcha and Rossky (2014***), which can describe the effective interaction between residues. The hydrophobicity value *λ*, ranging from 0 to 1, varies for different residues. The arithmetic average is used for the interaction parameter between two residues, i.e. hydrophobicity value *λ*_*i*,*j*_ = (*λ*_*i*_ + *λ*_*j*_)/2 and amino acid size *σ*_*i*,*j*_ = (*σ*_*i*_ + *σ*_*j*_)/2. The energy function of the short-range pairwise interaction is given by

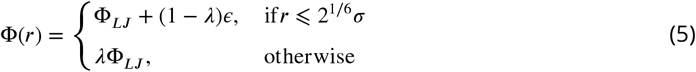

where Φ_*LJ*_ is standard 6-12 Lennard-Jones potential

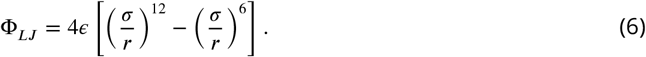

The free parameter *ϵ* is set as 0.2 kcal/mol, with which the molecular simulations can best reproduce the experimental results of *R*_*g*_ values for a set of IDPs.

### Simulation Framework

We used a slab simulation scheme in the coexistence simulations. Initially, 125 chains were randomly distributed in a cubic box with periodic boundary conditions at 200 K. The box length is first scaled to 15 nm, which can effectively prevent protein chains from interacting with their periodic images generated by periodic boundary conditions. Then, the *x*- and *y*- dimensions were held fixed and the *z*-dimension is extended to 280 nm. The temperature is then gradually increased to the target temperature and the simulations are conducted at different temperatures for 1 *μ*s in the NVT ensemble maintained by a Langevin thermostat, with a mass-normalized damping rate of 0.01 ps^−1^. A time step of 0.01 ps was used for all the simulations. We used HOOMD-Blue v2.9.7 packages ***Anderson et al. (2020***) and OpenMM v8.2.0 ***Eastman et al. (2023***) to conduct the simulations.

The slab density profile along the *z*-axis is then determined after the system had equilibrated. In order to avoid the influence of periodic boundary conditions, we first move the center of mass of the system along the *z*-axis direction to the position of *z* = 0. We then determined the distribution of dense phase and dilute phase of the phase separated system using the defined thresholds. The area with density higher than 0.95 of the maximum density (*ρ*_max_) is treated as dense phase, and their average density is regarded as the phase density of the dense phase (*ρ*_dense_). For the dilute phase, the threshold is taken as *ρ*_min_ + 50 mg/mL, where *ρ*_min_ is the minimum density. The density of dilute phase (*ρ*_dilute_) is then determined by the same method. If *ρ*_max_ − *ρ*_min_ < 50 mg/mL, the system is considered not to undergo phase separation.

### Critical Temperature

The critical temperature was determined based on the vapor-liquid interfacial properties of Lennard-Jones chains ***Blas et al. (2008***). The critical temperature *T*_*c*_ was obtained from *ρ*_dense_ and *ρ*_dilute_ at different temperatures by

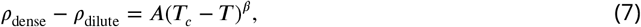

where *β* is the critical exponent, which was set to 0.325 ***Rowlinson and Widom (2013***) and *A* is the fitting parameter. Data with temperature below the critical temperature was used to fit this equation. The phase density at critical temperature was determined by the law of rectilinear diameters ***Davies (1912***)

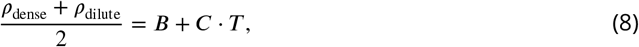

with *B* and *C* being the free parameters determined from the simulation data. At the critical temperature, the distinction between dense phase and dilute phase disappears, which means *ρ*_dense_ = *ρ*_dilute_ = *ρ*_*c*_. The phase density was then calculated by *ρ*_*c*_ = *B* + *C* ⋅ *T*_*c*_. The phase diagram was then constructed from the dense- and dilute-phase densities as functions of temperature.

### Bead-level second virial coefficient

The bead-level second virial coefficient was calculated from the effective pair potential as

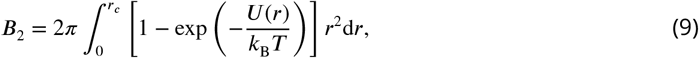

where *U* (*r*) is the desolvation-aware effective pair potential defined in Equation (1). The upper limit of integration *r*_*c*_ is set as 3*σ*, which is sufficiently large to capture the full range of interactions while ensuring numerical convergence. The reduced value *B*_2_/*σ*^3^ was used to compare the integrated effective attraction under different desolvation parameters. All *B*_2_ values reported in ***Figure 2—figure Supplement 1***G, H were evaluated at the same reduced temperature. This bead-level quantity provides a pair-potential-level measure of effective attraction.

### Temperature Normalization of the Generalized Diffusion Coefficient

To investigate the intrinsic influence of the interaction potential on chain dynamics, we must isolate the structural contributions from the trivial thermal acceleration. While keeping the quench depth (*T* /*T*_c_) constant provides a comparison at matched quench depth, it necessitates varying the absolute temperature *T*, which inherently alters the base diffusion rate via thermal fluctuations. To resolve this, we adopt the constant quench depth approach to ensure thermodynamic consistency and introduce a normalized metric to factor out the leading linear thermal contribution.

Motivated by the Einstein relation for Langevin diffusion, we describe the leading temperature dependence of the generalized diffusion coefficient *K* in terms of the thermal energy *k*_B_*T* and the effective friction coefficient *ζ* :

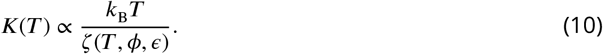

This approximation is motivated by the weakly subdiffusive dynamics observed here, for which *α* ≈ 0.91 and remains nearly constant across the simulated conditions. Here, the effective friction coefficient *ζ* encapsulates the resistance to chain motion. Following the hybrid theoretical framework proposed by Macedo and Litovitz ***Macedo and Litovitz (1965***), which combines free-volume theory with activated rate theory, we phenomenologically factorize the effective friction into packing-dependent and energy-barrier-dependent contributions:

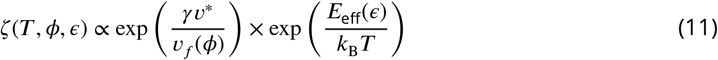

where the first term represents the hindrance due to limited free volume (*v*_*f*_), governed by the packing density *ϕ*, and the second term is the Arrhenius factor describing the activated hopping over an effective energy barrier *E*_eff_, which corresponds to the roughness of the energy landscape induced by the desolvation potential.

In our analysis, by fixing the quench depth (*T* /*T*_c_), we expect the macroscopic packing density *ϕ* to remain approximately comparable across different systems (as verified by the phase diagram in main text). Consequently, the free-volume contribution to friction is effectively invariant. Substituting this back into Equation (10), the diffusion coefficient scales as:

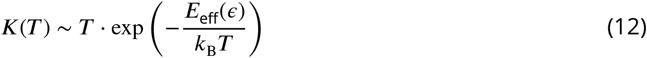

The prefactor *T* represents the leading linear thermal acceleration inherent to Langevin dynamics. To better isolate the intrinsic energy-landscape signature, we define the reduced diffusion coefficient 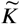 by normalizing out this explicit linear thermal contribution:

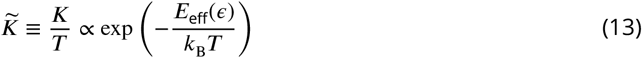

Thus, 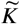 serves as a physically motivated metric to better isolate the intrinsic influence of the energy function on chain dynamics. By normalizing out the explicit linear thermal contribution inherent to Langevin dynamics, this definition helps decouple the specific kinetic modulation imposed by the desolvation potential from the background thermal acceleration and thermodynamic driving forces. Consequently, 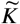 allows for a more direct assessment of how the interaction potential alters transport properties after removing the leading linear temperature effect required to maintain thermodynamic consistency.

### Calculation of Normalized Density Variance

To quantify the rate of density fluctuation growth during the early stage of phase separation, we first computed the instantaneous spatial variance of the density profile along the *z*-axis, 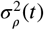, defined as

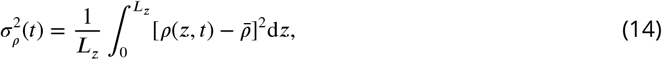

where *ρ*(*z, t*) is the local density at position *z* and time *t*, and 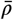 is the mean density of the system. To enable a fair comparison of kinetics across systems with distinct interaction parameters (*ϵ*_ss_ and *ϵ*_b_), it is crucial to decouple the rate of phase separation from the difference in equilibrium dense-phase densities. Therefore, we normalized the instantaneous variance by its equilibrium baseline:

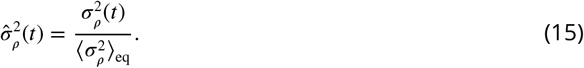

Here, the normalization factor 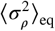 is the time-averaged density variance calculated from the final 30% of the simulation process. This period corresponds to the steady state characterized by the formation of a well-defined slab with stabilized phase densities. By using this normalized metric 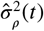, we effectively factor out the variations in final density contrast, allowing the observed differences in the time-evolution curves to more directly reflect the acceleration or deceleration of the kinetic process itself.

### Desolvation Parameters

Because residue-pair-specific experimental desolvation data were unavailable, the empirical parameters *α*_b_ and *α*_ss_ were estimated using all-atom molecular dynamics simulations of amino acid analogues with explicit solvent. The potential of mean force (PMF) obtained from the simulation results is shown in ***Figure 1—figure Supplement 1***. Methane (CH_4_), methanol (CH_3_OH), acetate ion 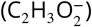, ammonium ion 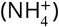, and methanamide (CH_3_NO) were chosen as representative amino acid analogues for non-polar, polar, negatively charged, positively charged, and backbone-like residues, respectively. For each analogue pair, *α*_b_ and *α*_ss_ were obtained by fitting the non-bonded energy function with desolvation effects (Equation (1)) to the simulation-derived PMF. The *λ* values assigned to the analogues were taken from the corresponding amino acids. Specifically, we used *λ*_GLY_ = 0.649 for methane, *λ*_SER_ = 0.595 for methanol and *λ*_CYS_ = 0.595 for methanamide. The results are shown in Table 1.

## Data Availability

All simulation codes, input files, and scripts for plotting figures of the manuscript are openly available on GitHub at https://github.com/kaizhangnju/desolvation-CG-model. Additional data supporting the findings of this work are available from the corresponding author upon reasonable request due to large file size constraints.

## Acknowledgments

This work was supported by the Fundamental and Interdisciplinary Disciplines Breakthrough Plan of the Ministry of Education of China (JYB2025XDXM502), the National Natural Science Foundation of China (12574224 and 12347102), the Basic Research Program of Jiangsu Province (BK20253050), and the Wenzhou Institute, University of Chinese Academy of Sciences (WIUCASQD2021010). The authors also acknowledge support from Nanjing Kunpeng&Ascend Center of Cultivation, the HPC Center of Nanjing University, the e-Science Center of Nanjing University, the Nanjing Key Laboratory for Cardiovascular Information and Health Engineering Medicine (funded by the Nanjing Municipal Health Commission), and its Jiangsu counterpart.

---

**1—figure supplement 1.**
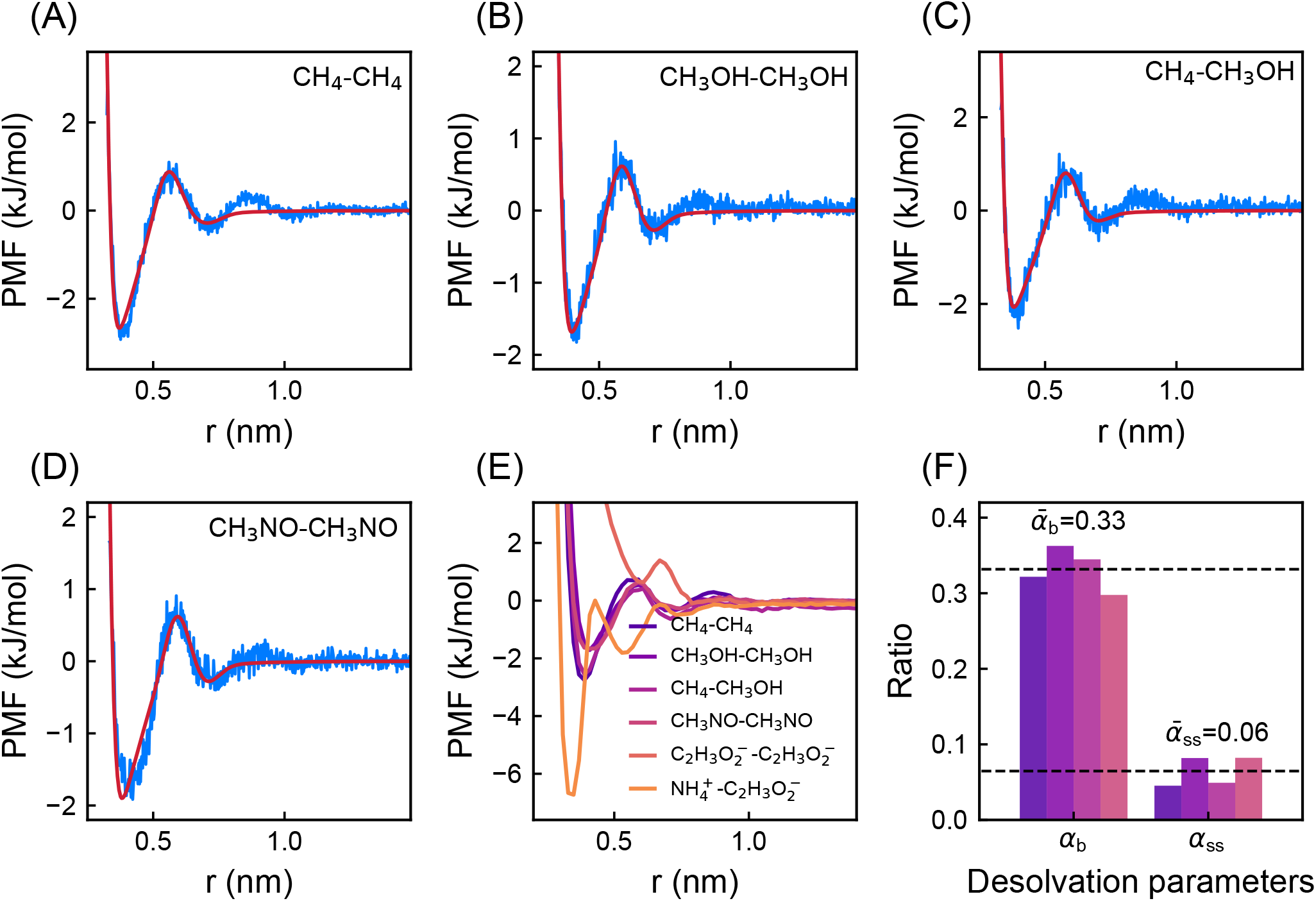
Desolvation-related PMF features and fitted parameters from all-atom analogue simulations. (A-D) PMFs obtained from all-atom simulations of representative amino-acid analogue pairs, together with fits using the desolvation-aware effective potential in Equation 1. (E) Overlay of fitted PMF profiles for different analogue pairs, illustrating the shared double-minimum/barrier structure and the pair-dependent variation in desolvation features. (F) Desolvation coefficients *α*_b_ and *α*_ss_ obtained by expressing the fitted barrier height and solvent-separated minimum depth relative to the global reference energy scale *ϵ* used in the effective potential.

**2—figure supplement 1.**
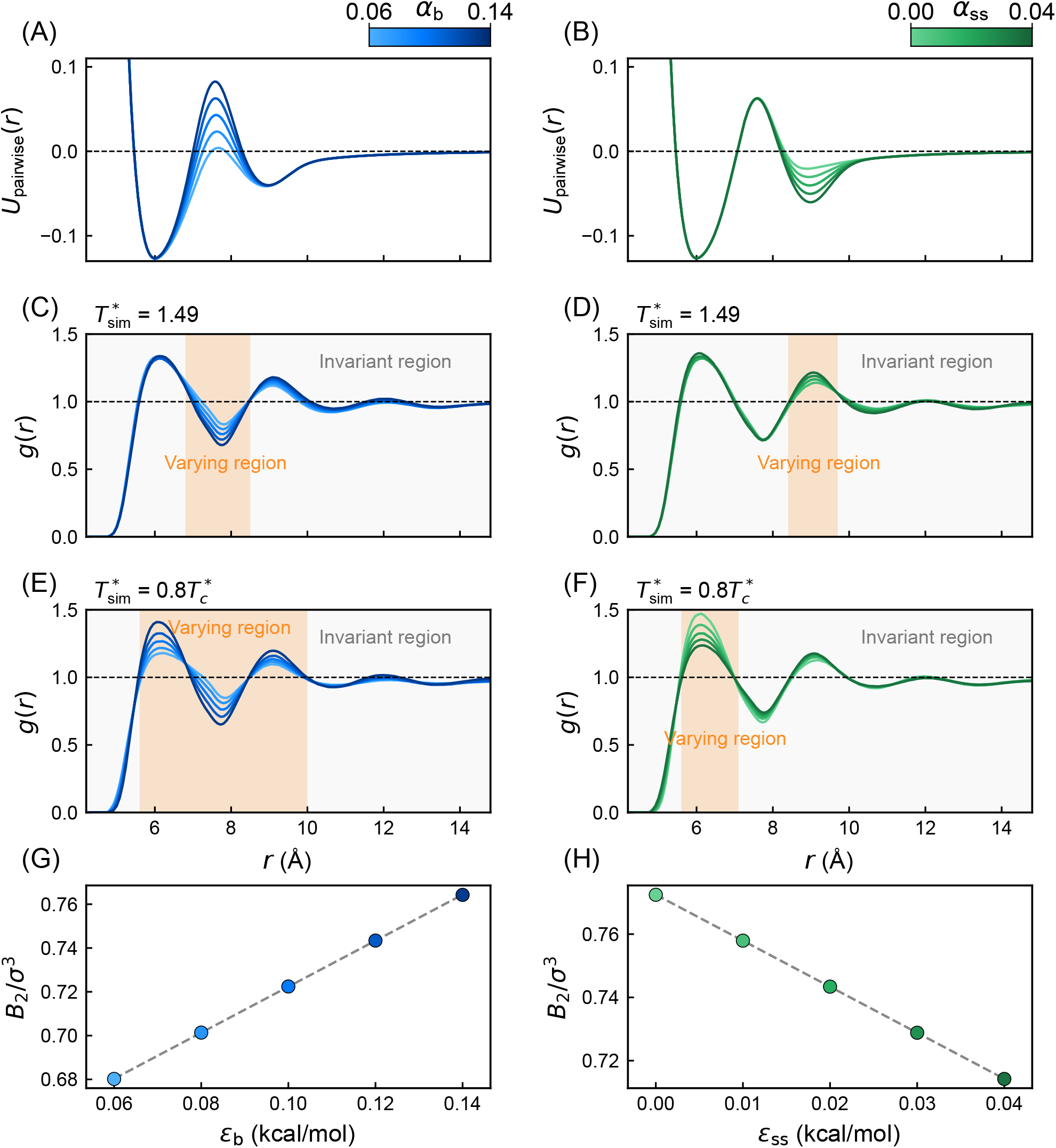
Microscopic residue-pair distributions and integrated effective attraction under varying desolvation parameters. (A, B) Schematic illustration of the effective pair potential corresponding to different values of *ϵ*_b_ or *ϵ*_ss_. (C, D) Radial distribution functions of all residue pairs under varying *ϵ*_b_ (C) or *ϵ*_ss_ (D) at the same reduced temperature, *T* ^∗^ = 1.49. (E, F) Radial distribution functions of all residue pairs in the dense phase at the same normalized temperature 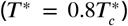. (G, H) Bead-level second virial coefficient (*B*_2_/*σ*^3^) calculated from the effective pair potential under varying *ϵ*_b_ at fixed *ϵ*_ss_ = 0.02 kcal/mol (G) and varying *ϵ*_ss_ at fixed *ϵ*_b_ = 0.12 kcal/mol (H).

**5—figure supplement 1.**
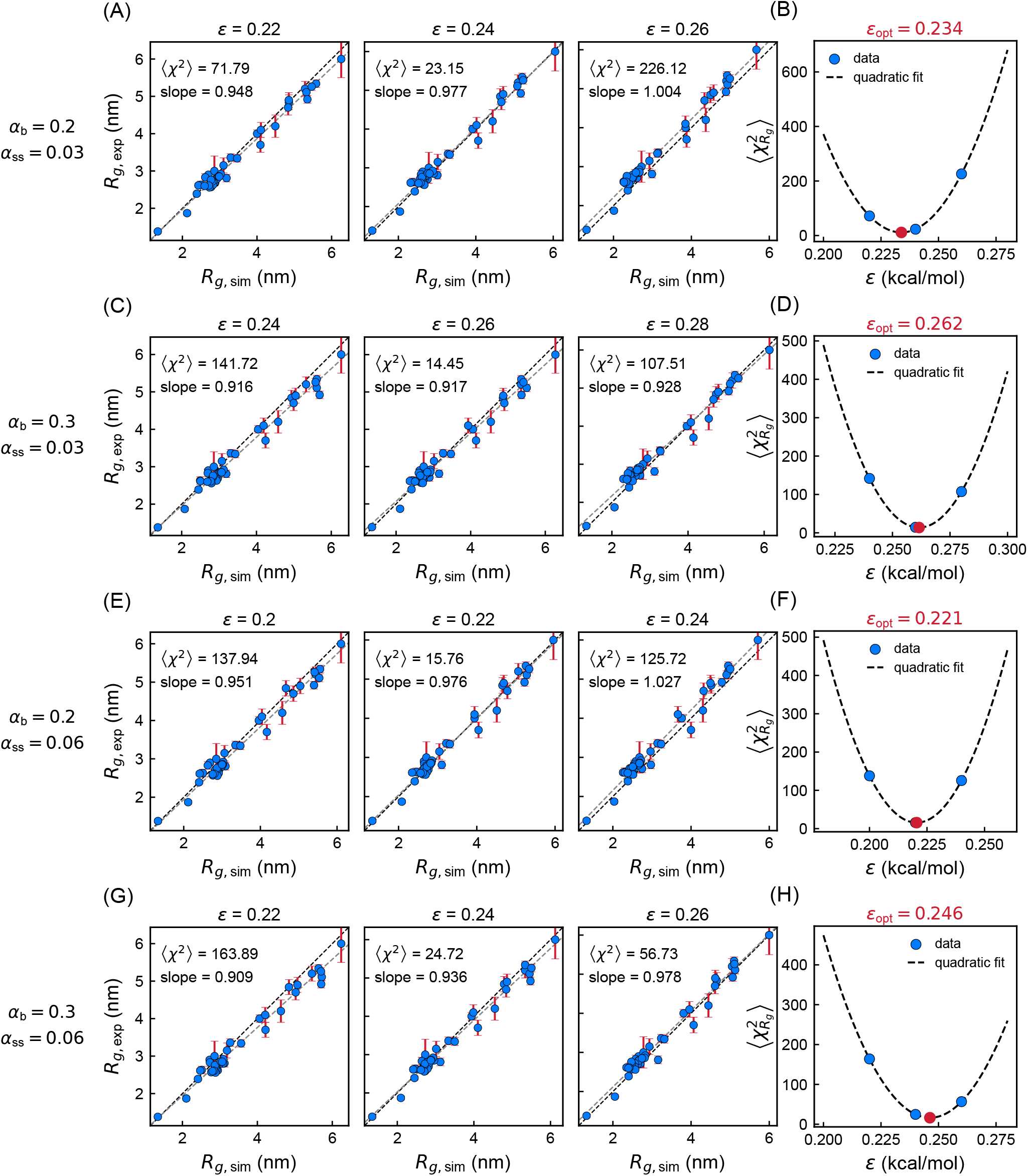
Optimization of the CALVADOS2 + desolvation energy scale using experimental *R*_*g*_ data. (A, C, E, G) Correlations between experimental and simulated *R*_*g*_ values for the IDP dataset under selected desolvation-parameter combinations. Each panel corresponds to simulations performed at fixed *α*_b_ and *α*_ss_ with different values of the overall energy scale *ϵ*. (B, D, F, H) Corresponding 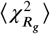 values as a function of *ϵ*, used to identify the optimal energy scale for each desolvation-parameter set. The optimized *ϵ* values are indicated in each panel.

**5—figure supplement 2.**
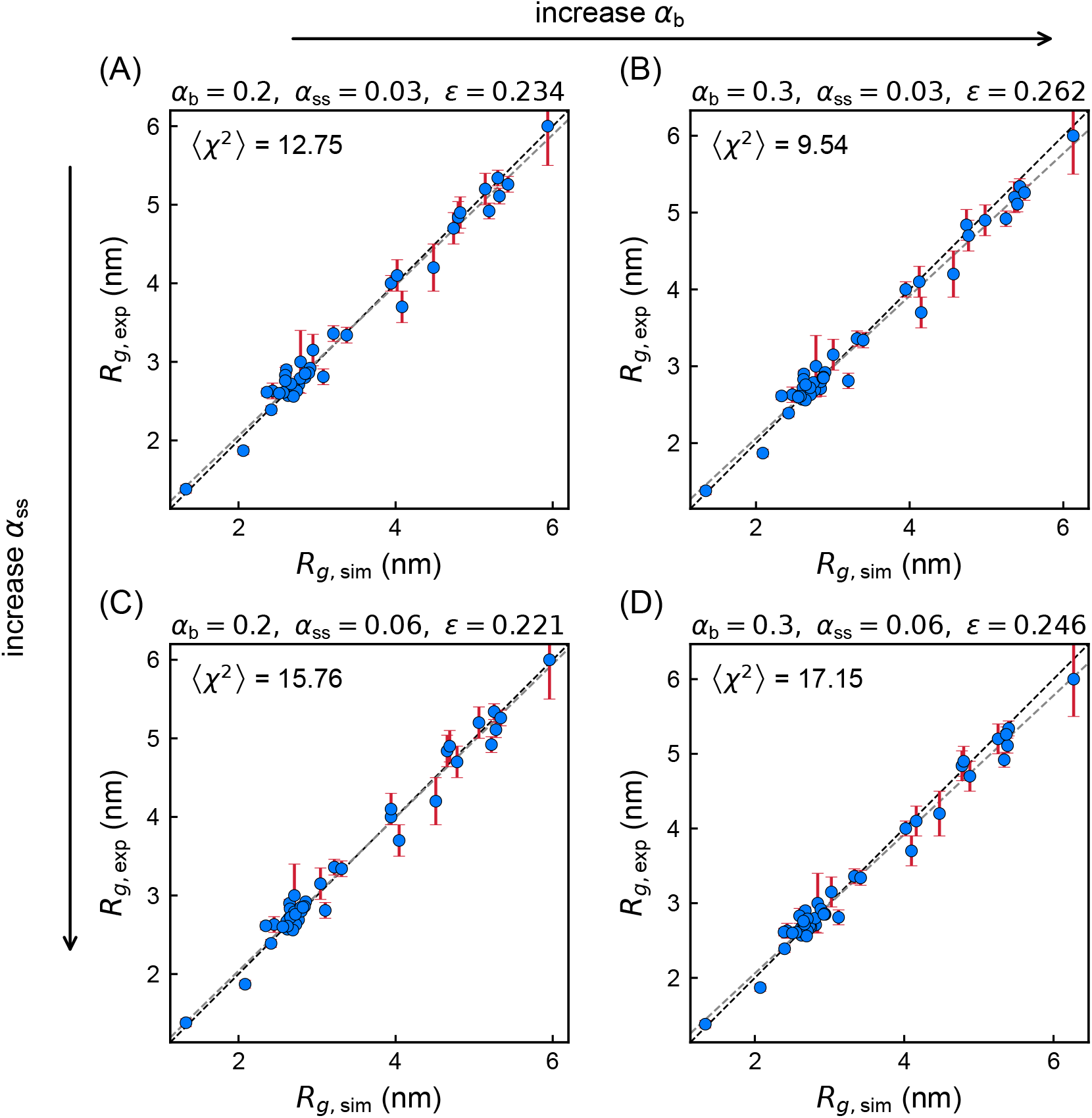
Global scan of CALVADOS2 + desolvation parameter combinations against experimental *R*_*g*_ values. Comparison between experimental and simulated *R*_*g*_ values for different combinations of *α*_b_ and *α*_ss_, with the overall energy scale (*ϵ*) optimized separately for each parameter set. The corresponding optimized (*ϵ*) values are shown to illustrate how the energy rescaling compensates for different desolvation strengths while preserving agreement with single-chain dimensions.

**5—figure supplement 3.**
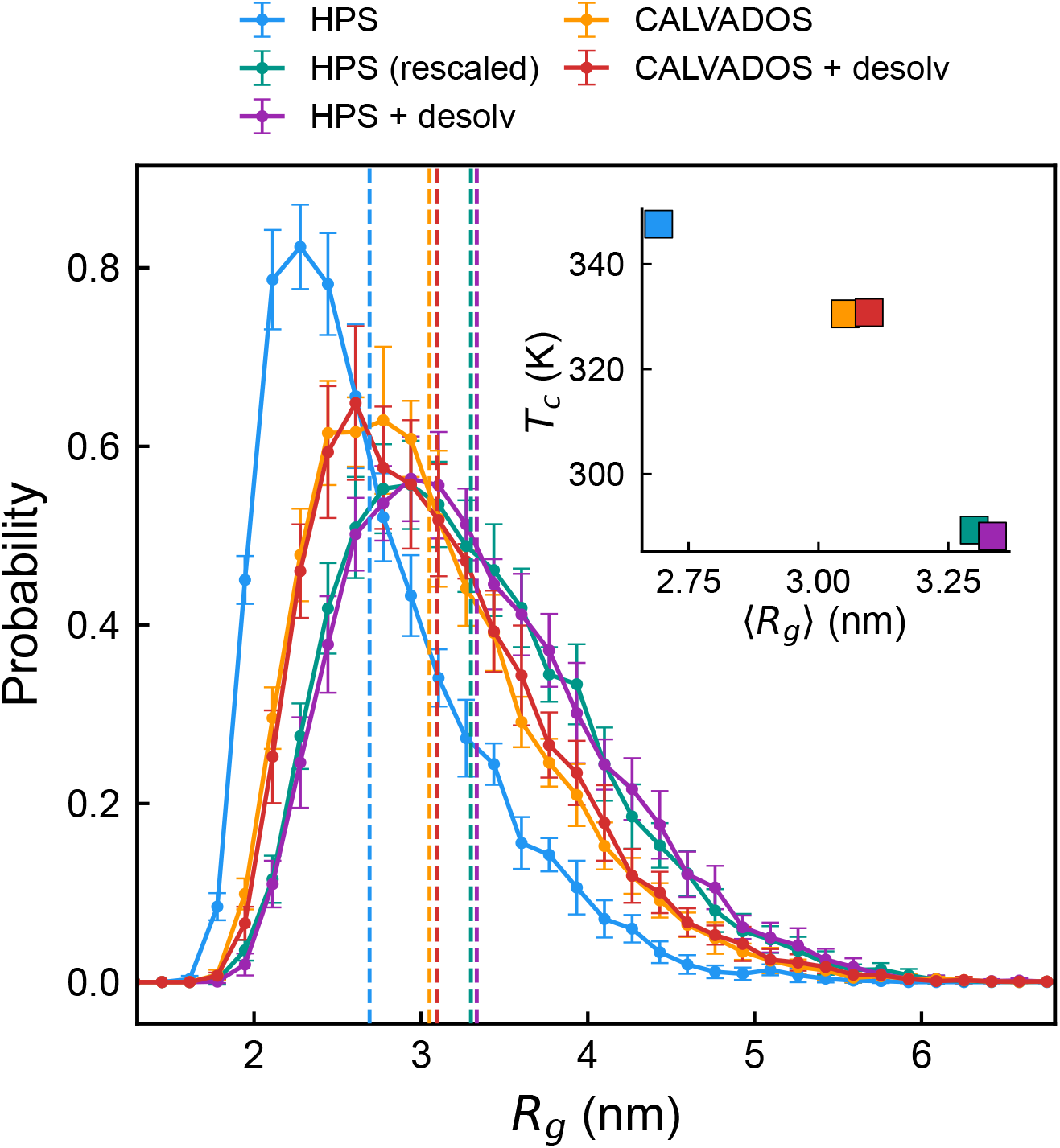
Single-chain *R*_*g*_ comparison of FUS LC under different coarse-grained model variants. Single-chain radius of gyration of FUS LC simulated using the original HPS model, HPS + desolvation, energy-rescaled HPS, original CALVADOS2, and CALVADOS2 + desolvation models. The energy-rescaled HPS model was included to separate the effect of global interaction-strength rescaling from the effect of adding desolvation terms.

**5—figure supplement 4.**
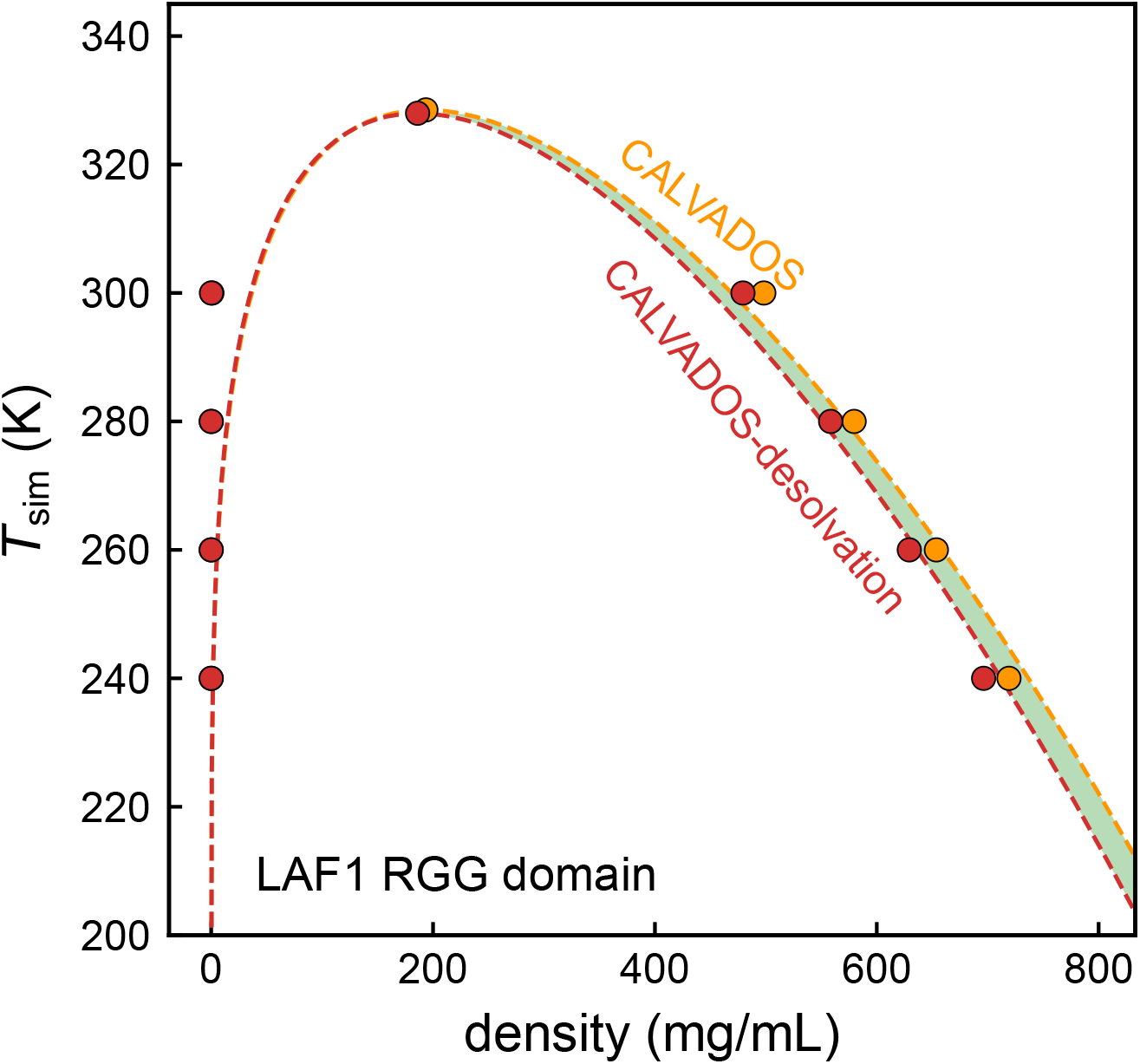
Phase behavior of the LAF-1 RGG domain simulated with CALVA-DOS2 and CALVADOS2 + desolvation. Coexistence curves of the LAF-1 RGG domain simulated using the original CALVADOS2 model and the desolvation-aware CALVADOS2 model. Symbols show the coexisting dilute- and dense-phase densities obtained from slab simulations at different simulation temperatures, and dashed curves represent binodal fits using the same critical-scaling procedure as in the main text.

